# Interpreting the pathogenicity of Joubert Syndrome missense variants in *Caenorhabditis elegans*

**DOI:** 10.1101/2020.05.22.110668

**Authors:** Karen I. Lange, Sofia Tsiropoulou, Katarzyna Kucharska, Oliver E. Blacque

## Abstract

Ciliopathies are inherited disorders caused by cilia defects. Variants in ciliopathy genes are frequently pleiotropic and represent excellent case studies for interrogating genotype-phenotype correlation. We have employed *Caenorhabditis elegans* and gene editing to characterise two pathogenic biallelic missense variants (P74S, G155S) in B9D2/*mksr-2* associated with Joubert Syndrome (JBTS). B9D2 functions within the MKS module at the transition zone (TZ) ciliary subcompartment, and regulates the cilium’s molecular composition and signaling function. Quantitative assays of cilium/TZ structure and function, together with knock-in reporters, confirm both variant alleles are pathogenic. G155S causes a more severe overall phenotype and disrupts endogenous MKSR-2 organisation at the TZ. Recapitulation of the patient biallelic genotype shows that heterozygous worms phenocopy worms homozygous for P74S. This study also reveals a close functional association between the B9 complex and TMEM216/MKS-2. These data establish *C. elegans* as a paradigm for interpreting JBTS mutations, and provide insight into MKS module organisation.

## INTRODUCTION

Ciliopathies represent at least 35 genetically inherited disorders, and more than 200 associated genes, that collectively affect the development and/or homeostasis of many body tissues and organs. Examples include Joubert syndrome (JBTS), Meckel syndrome (MKS), and nephronophthisis (NPHP), which present with overlapping clinical presentations including brain malformations, bone abnormalities, sensory defects, left-right asymmetries, cystic kidney disease, retinal degeneration, and cognitive impairment (Wheway et al., 2019). Ciliopathies result from defects in motile or primary cilia that extend from the surfaces of most eukaryotic cell types. Non-motile primary cilia function as antenna-like organelles, responding to many extracellular sensory cues such as light, odorants and osmotic strength, as well as cell-cell signaling molecules (eg. sonic hedgehog) that regulate cell behaviour, tissue formation and homeostasis (Anvarian et al., 2019). The canonical primary cilium consists of a cylinder of 9 doublet microtubules extending from a plasma membrane-anchored basal body (Satir et al., 2010).

Despite the cilium’s cytosol and membrane being contiguous with those of the cell, the organelle is highly compartmentalised with a unique molecular composition and signaling environment. Compartmentalisation is achieved by active transport pathways (eg intraflagellar transport) that bring proteins into and out of the cilium, as well as gating mechanisms at the ciliary base (Jensen & Leroux, 2017; Nachury & Mick, 2019). A major component of the ciliary gate is the proximal-most 0.2-1 μm of the axoneme, termed the transition zone (TZ). Characterised by Y-links that connect the ciliary membrane and axonemal microtubules, the TZ is thought to serve as a membrane and size-dependent cytosolic diffusion barrier to protein exchange between the cilium and the main cell body (Garcia-Gonzalo & Reiter, 2017; Reiter et al., 2012). At least 28 ciliopathy proteins, many associated with JBTS, MKS and NPHP, localise at the TZ, functioning within multisubunit functional modules and complexes to regulate ciliary gating and signaling (Garcia-Gonzalo & Reiter, 2017; Reiter & Leroux, 2017). Mutations in TZ proteins disrupt the ciliary localisation of important signalling molecules such as the Shh pathway regulator, Smoothened (Garcia-Gonzalo et al., 2011; Shimada et al., 2017; Shi et al., 2017).

TZ structure, function and molecular organisation is especially well understood in *Caenorhabditis elegans*, where cilia are found at the ends of the dendrites of 60 sensory neurons (Inglis et al., 2007). Many ciliopathy proteins and associated pathways are conserved in the worm (Kim et al., 2018; van Dam et al., 2019), and *C. elegans* studies have led to seminal discoveries that have greatly instructed our understanding of cilia biology in vertebrates and mammals (Blacque & Sanders, 2014; Ganner & Neumann-Haefelin, 2017; Mok & Héon, 2012; Molinari & Sayer, 2020). Genetic studies in *C. elegans* have revealed that TZ proteins function within two distinct entities termed the ‘NPHP’ and ‘MKS’ modules (Williams et al., 2008, 2011). The NPHP module is composed of NPHP-4/Nphp4 and NPHP-1/Nphp1 (Winkelbauer et al., 2005). The much larger MKS module is subdivided into core (MKS-1/MKS1, MKSR-1/B9D1, MKSR-2/B9D2, MKS-2/TMEM216, and TMEM-231/TMEM231), intermediate (TMEM-107/TMEM107), and peripheral (MKS-3/TMEM67, MKS-6/CC2D2A, JBTS-14/TMEM237, TMEM-17/TMEM17, and TMEM-218/TMEM218) submodules, which assemble at the TZ in a hierarchical fashion (core-intermediate-peripheral) (Bialas et al., 2009; Huang et al., 2011; Lambacher et al., 2016; C. Li et al., 2016; Roberson et al., 2015; Williams et al., 2008, 2010, 2011). The TZ recruitment of the entire MKS module depends on CEP-290/Cep290 (C. Li et al., 2016; Schouteden et al., 2015). Additionally, MKS-5/RPGRIP1L recruits the NPHP *and* MKS modules, *and* CEP-290, to the TZ, and is therefore the master regulator of TZ assembly in *C. elegans* (Jensen et al., 2015). Loss of individual MKS or NPHP module proteins results in abnormal ‘leakage’ of membrane proteins into and out of sensory cilia due to disruption of the TZ-associated membrane diffusion barrier (Cevik et al., 2013; Williams et al., 2011). However, with regard to ciliogenesis and TZ formation, the NPHP and MKS modules function redundantly, with defects only observed when a gene from both modules is disrupted (Williams et al., 2008, 2011). While the hierarchical blueprint of TZ organisation is largely conserved in higher organisms there are some differences. Most notably, in mice, CEP290 recruits the NPHP module, and Rpgrip1/Rpgrip1l act synergistically to recruit the MKS module (Wiegering et al., 2018). Tissue specific differences have also been reported for TZ component Mks6 in mice (Lewis et al., 2019).

Since many ciliopathy genes are pleiotropic and frequently show high levels of variability in their phenotypic expressivity, ciliopathy disorders are an excellent paradigm for interrogating human genotype-phenotype correlation. Despite this, most cell and animal model studies have traditionally studied ciliopathy mechanisms using null allele, gene depletion, or complementation-based approaches. Whilst invaluable for revealing important aspects of disease mechanism, these approaches do not interrogate the phenotypic effects of specific variants as they occur in patients. Indeed, genetic compensation arising from a gene knockout is now a well recognised phenomenon (El-Brolosy et al., 2019). In this study, we have explored the use of *C. elegans* to model missense mutations associated with a ciliopathy. Specifically, we focussed on biallelic mutations (P74S, G155S) in B9D2, previously reported in a Joubert syndrome (JBTS) patient with a molar tooth sign, polydactyly, seizures, and a cleft palate (R. Bachmann-Gagescu et al., 2015). The effect of these mutations on B9D2 function is not known.

B9D2 is one of three MKS module proteins with a B9 domain, which is a subfamily of the C2 phospholipid binding domain (Lemmon, 2008) found only in ciliated eukaryotes (Zhang & Aravind, 2010). These three proteins (MKS1, B9D1 and B9D2) are proposed to form a complex (Boldt et al., 2016; Chih et al., 2011; Garcia-Gonzalo et al., 2011; Gupta et al., 2015; S. Li et al., 2004; Simonis et al., 2009; Zhao & Malicki, 2011). B9D2 is required for ciliogenesis in many organisms including mice (Town et al., 2008), paramecium (Ponsard et al., 2007) and loss of B9D2 is associated with developmental defects in zebrafish (Dowdle et al., 2011). As in *C. elegans*, the B9 complex has been shown to serve ciliary gating functions in *Drosophila* and mice (Basiri et al., 2014; Chih et al., 2011). Mutations in the B9 complex have been linked to various ciliopathies including JBTS (R. Bachmann-Gagescu et al., 2015; Bader et al., 2016; Romani et al., 2014; Slaats et al., 2016) and MKS (Cui et al., 2011; Dowdle et al., 2011).

Here, we used CRISPR-Cas9 genome editing to generate *C. elegans* alleles carrying the P74S and G155S mutations in the *C. elegans* B9D2 (*mksr-2*). Via a knock-in tag, we found that both endogenously-expressed variant proteins localised to the TZ with reduced efficiency, while the G155S variant displaying dramatically altered TZ distribution when visualised using super-resolution microscopy. Using quantitative assays of cilium structure and function, as well as ultrastructural analyses, we identified a less severe overall recessive phenotype for P74S-containing worms, compared with worms harbouring G155S, whose phenotype is nearly as severe as a reference null allele. Consistent with this observation, an assay for the TZ membrane diffusion barrier revealed that an RPI-2::GFP reporter leaked into cilia of worms with the G155S, but not the P74S, mutation. Unlike the reference null, endogenous MKS module protein reporters still assemble at the TZ of the *mksr-2* mutants, although the TZ level and distribution of these reporters is disrupted, especially for MKS-2. We also found that endogenous MKS module reporters do not recapitulate several previously reported TZ localisation phenotypes. Lastly, compound heterozygous worms (P74S/G155S) that reflect the biallelic genotype of the JBTS patient display a phenotype that approaches that of worms recessive for the P74S mutation. Taken together these data demonstrate the utility of *C. elegans* for interpreting the pathogenicity of missense alleles in JBTS. Our work also uncovers new insight into MKS module organisation at the TZ, including a close functional association between the B9 complex and TMEM216/MKS-2.

## RESULTS

### Modelling pathogenic missense B9D2 variants associated with Joubert Syndrome

The B9D2 protein is highly conserved from vertebrates to the nematode *C. elegans* (**Figure 1A**). Human B9D2 and the *C. elegans* MKSR-2 orthologue possess 63.4% amino acid similarity (100% length, BLAST e-value 3e-52). The protein is composed of a B9 domain (aa 2-118) and a C-terminal region with no predicted domains. B9 domains are found only in ciliated eukaryotes and represent a subfamily of lipid and membrane-associating C2 domains (Zhang & Aravind, 2010); the function of the C-terminal region of B9D2 is unknown.

**Figure 1.**
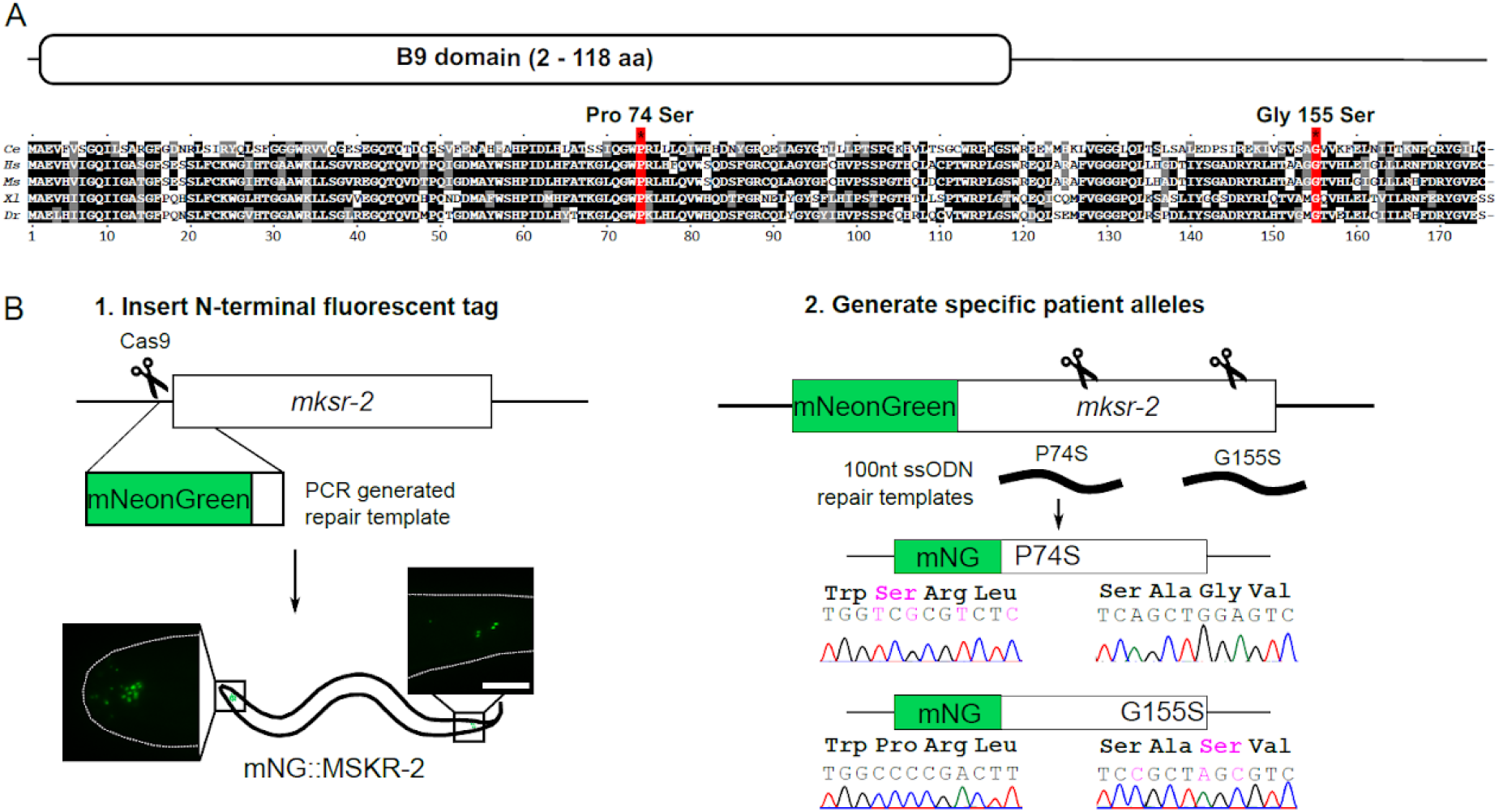
CRISPR-Cas9 engineering of P74S and G155S into the orthologous positions of *C. elegans* mksr-2. **(A)** Protein alignment of B9D2 sequences from *C. elegans* (Q9N423), *H. sapiens* (Q9BPU9), *M. musculus* (Q3UK10), *X. laevis* (Q6GN70), and *D. rerio* (Q6DGZ1). UniportKB accession numbers are indicated. B9D2 proteins contain a B9 domain (amino acids 2-118) and a C-terminal tail with no predicted domains. Two pathogenic Joubert Syndrome mutations (red), Pro74Ser and Gly155Ser, affect conserved residues. Protein sequences were aligned in Clustal Omega and BoxShade 3.21 was used to generate the figure. **(B)** CRISPR/Cas9 genome editing was used to introduce a fluorescent mNeonGreen (mNG) tag to the N-terminus of the endogenous *mksr-2* gene. Scale bar; 10 μm. The mNG::MKSR-2 tagged allele was then used to engineer two pathogenic missense variants, Pro74Ser and Gly155Ser, associated with Joubert Syndrome. Silent and missense engineered mutations are shown in magenta. ssODN; single-stranded oligonucleotide. Detailed information about the CRISPR approach is in Figure S1.

Since the B9D2 protein is highly conserved, we reasoned that *C. elegans* would be an excellent organism to model pathogenic patient variants of this protein in a biological system. Specifically, we focussed on two missense variants: Pro74Ser in the B9 domain and Gly155Ser in the C-terminal region (**Figure 1A**). Both mutations are reported to be pathogenic in a biallelic patient with JBTS (R. Bachmann-Gagescu et al., 2015). The compound heterozygosity of P74S and G155S represents an excellent paradigm for interpreting the individual and combined effects of variants at one locus. Patient allele phenotypes will be compared to those of a null deletion allele, *mksr-2(tm2452)* (hereafter termed *mksr-2(Δ)*) (Bialas et al., 2009; Williams et al., 2008, 2011).

To facilitate assessment of endogenous protein localisation, prior to generating the patient alleles, we used CRISPR-Cas9 to knock-in *C. elegans* optimised mNeonGreen (mNG) (Hostettler et al., 2017) with a short 7aa flexible linker (GTGGGGS) at the 5’ end of the *mksr-2* locus (**Figure 1B**). Specifically, a crRNA was designed to target a PAM site near the start of the gene and a PCR product containing the mNG sequence with 35bp homology arms was used as the repair template (Table S1). Germline editing was performed via a co-CRISPR strategy, involving pre-assembled CRISPR-Cas9 ribonucleoprotein complexes injected into the gonad of young adult hermaphrodite worms (Paix et al., 2015). This generated the *mksr-2(oq108)* allele which we refer to as *mNG::mksr-2(+)*.

Next, we used co-CRISPR with 100 nucleotide single stranded repair templates to engineer the P74S and G155S variants into the *mNG::mksr-2(+)* strain (Paix et al., 2017). Both variants were generated in a single CRISPR-Cas9 injection (**Figure 1B**). PAM sites near each locus were targeted, and restriction enzyme cut sites introduced via the repair templates to facilitate variant detection (**Figure S1**). Sanger sequencing of *mksr-2* confirmed the correct incorporation of the variant, and the absence of undesired mutations in the gene. We isolated one allele for each variant: *mksr-2(oq108,oq125)* (*mNG::mksr-2(P74S)*) and *mksr-2(oq108,oq126)* (*mNG::mksr-2(G155S)*). Prior to phenotypic analyses, strains were out-crossed twice with wild-type worms to remove any potential off target mutations generated by CRISPR-Cas9.

Finally, we also generated non-tagged versions of the variants: *mksr-2(oq137)* (*mksr-2(P74S)*) and *mksr-2(oq138)* (*mksr-2(G155S)*) using a modified PCR-based detection protocol, which is faster and less expensive than that using restriction enzymes (**Figure S1**). Briefly, engineered variants are detected using oligonucleotides that contain the silent and missense mutations; a PCR product is only amplified when the variants have been incorporated into the genome. More than one mutant primers can be used in a single reaction to simultaneously detect multiple variants.

### MKSR-2 distribution at the transition zone is severely altered by the G155S mutation

First, we investigated if the P74S and G155S mutations affect the ciliary localisation of endogenous MKSR-2. Analysis of the *mNG::mksr-2(+)* strain revealed that wild type MKSR-2 is localised almost exclusively at the TZs of cilia in the head (labial and amphid) and tail (PQR and phasmid) of the worm (**Figure 2A; S2**). Previous reports of MKSR-2 localisation in *C. elegans*, employing an overexpressed transgene, showed that MKSR-2::YFP localises at the TZ and in the more proximal periciliary region at the dendritic ending (Williams et al., 2008). Since the endogenously expressed mNG::MKSR-2 shows no such periciliary signal, we conclude that this previously reported pool of MKSR-2 is an artefact of overexpression.

**Figure 2.**
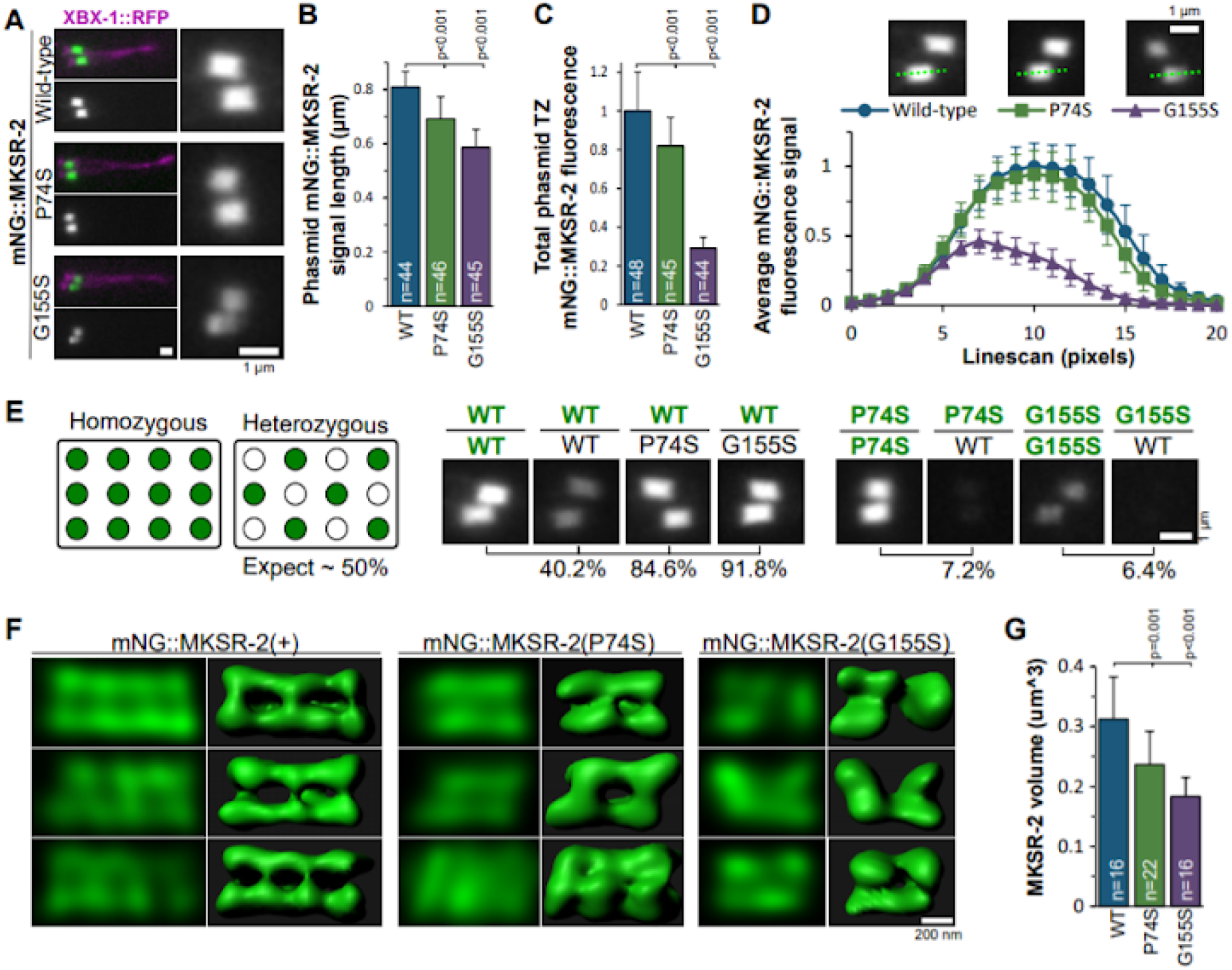
P74S and G155S mutations differentially disrupt the recruitment and organisation of endogenous MKSR-2 at the transition zone (TZ). **(A)** Representative images of one pair of phasmid cilia from worms expressing XBX-1::RFP (marks entire cilium) and endogenously-tagged mNG::MKSR-2 (WT and variants). A magnified view of the TZs (side on views) are shown in right-hand panels. Anterior is to the left. Scale bars; 1 μm. **(B)** Quantification of mNG::MKSR-2 signal length by calculating the “full width at half max” (FWHM) from a line scan down the length of the phasmid TZ. Data represents mean with S.D. p-values from unpaired t-tests. N; total number of TZs measured. **(C)** Quantification of the total fluorescence intensity of mNG::MKSR-2 at the TZ of phasmid cilia. Background fluorescence was subtracted and the signal normalised to wild-type. Data represents mean and S.D. p-values from unpaired t-tests. N; number of phasmid pairs measured. **(D)** Average fluorescence intensity from line scans (20 pixels = 1.6 μm) of phasmid cilia showing the distribution of mNG::MKSR-2 along the TZ length. Green dotted lines in images show TZ line scan examples. Error bars represent S.D. n=12 for each dataset. **(E)** Assessment of the efficiency of endogenous mNG::MKSR-2 recruitment to the phasmid TZ using strains heterozygous for the mNG::MKSR-2 alleles. In heterozygous worms, if mNeonGreen and non-tagged MKSR-2 alleles are recruited to the TZ with the same efficiency, the fluorescence intensity should be 50% of what is observed for worms homozygous for the tagged allele. Representative images are from a single plane. Relative % fluorescence is indicated for each heterozygous genotype, compared to the homozygous reference (set at 100%). Green lettering indicates an mNG-tagged allele of *mksr-2*. **(F)** Representative super-resolution confocal images of endogenous mNG::MKSR-2 in phasmid cilia. A single slice from the z-stack and corresponding 3D model are shown. Scale bars; 200 nm. All images are oriented with the proximal end of the TZ to the left. **(G)** Quantification of the volume of the mNG::MKSR-2 compartment at the TZ of phasmid cilia. Data represents mean and S.D. p-values from unpaired t-tests. N; number of TZ measured.

Analysis of the *mNG::mksr-2(P74S)* and *mNG::mksr-2(G155S)* strains revealed near exclusive localisation of the MKSR-2 variants at the TZ, similar to the wild-type reporter (**Figure 2A**). However, compared with mNG::MKSR-2(+), the length of the TZ signal is 15-25% shorter for both variants (**Figure 2B**). In addition, the overall levels of the mNG::MKSR-2 variants at the TZ are moderately (P74S) or severely (G155S) reduced (**Figure 2C**). Furthermore, linescan analysis of the fluorescent signals revealed that the mNG::MKSR-2(G155S) protein is not homogeneously distributed at the TZ like mNG::MKSR-2(+) and mNG::MKSR-2(P74S). Instead, the TZ fluorescence signal for the G155S variant peaks at the proximal end and is lower at the distal end (**Figure 2D**; see also **Figure 2A**).

We investigated if the variants affected the efficiency of MKSR-2 targeting to the TZ by examining their endogenous localisations when a wild-type copy of the gene is present. Genetic crosses were used to generate heterozygous worms (**Figure S3A**). If the tagged protein is targeted to the TZ as efficiently as the untagged wild type protein, then the fluorescent TZ signal should be half that observed in homozygous worms (**Figure 2E**). Indeed, heterozygous mNG::MKSR-2(+)/+ worms show ~60% reduction in TZ fluorescence, indicating that mNG::MKSR-2(+) retains a near wild-type ability to localise at the TZ (**Figure 2E; S3B**). However, in worms heterozygous for mNG::MKSR-2(+) with either untagged MKSR-2(P74S) or MKSR-2(G155S), fluorescence is only decreased by 10-15% indicating that the untagged mutant proteins are unable to compete with wild-type MKSR-2 for TZ incorporation (**Figure 2E; S3B**). In support of this conclusion, the reverse experiment using mNG::MKSR-2(P74S)/+ and mNG::MKSR-2(G155S)/+ heterozygotes showed an almost complete loss of TZ localisation for the tagged variants when an untagged copy of wild-type MKSR-2 is present (**Figure 2E, S3B**). These data indicate that both the P74S and G155S drastically decrease the efficiency of MKSR-2 recruitment to the TZ.

To further assess the altered distribution of the endogenous MKSR-2 variants at the TZ we employed a scanning confocal-based super resolution microscopy methodology (Lam et al., 2017). With confocal imaging using a reduced pin hole size and deconvolution we obtained an optical resolution of approximately 150 nm. Three-dimensional models of the phasmid TZ, in axial (longitudinal) view, were constructed from z-stacks. Using this approach it was found that mNG::MKSR-2(+) is excluded from the inner core of the TZ and displays a periodic pattern at the outer (peripheral) region of the compartment, consisting of 3 or 4 peaks of axial signal spaced 243.6 nm apart (sd 46.9 nm, n=16) (**Figure 2F**). In some images, the signals appear to wrap around the TZ, possibly as a spiral (**Figure S4**). This distribution pattern for mNG::MKSR-2(+) is similar to what we previously described using STED microscopy for overexpressed reporters of other *C. elegans* MKS module proteins (Lambacher et al., 2016).

Consistent with the widefield imaging data, super-resolution imaging reveals a highly disrupted localisation for mNG::MKSR-2(G155S) (**Figure 2F**); typically, the protein is discontinuous along the TZ length, with 88% of analysed TZs (n=16) showing only 2 peaks of axial signal that are 357.2 nm apart (sd 141.2 nm, n=14, p=0.011), and an apparent loss of the spiral pattern. In addition, the ‘hollow TZ core’ is not always observed in the central plane of the images, and the 3D models show that mNG::MKSR-2(G155S) occupies a reduced overall TZ volume (**Figure 2F, G**). The super-resolution imaging also provided evidence of subtle TZ distribution defects for mNG::MKSR-2(P74S), which displays only 2-3 peaks of axial signal and a less pronounced spiral pattern; the reporter also occupies a modestly reduced TZ volume (**Figure 2F, G**). The decrease in axial peak number for mNG::MKSR-2(P74S) is likely due to the overall shortening of the fluorescence signal since the peaks are 262.5 nm apart (sd 47.3, n=19) which is similar to the wild-type reporter.

Overall, these data indicate that both P74S and G155S affect the TZ targeting and distribution of endogenous MKSR-2. While both alleles are pathogenic, the G155S variant has a more severe effect on the protein organisation at the TZ. The P74S amino acid substitution is found in the B9 domain but does not appear to abolish association with the membrane since the hollow core is still observed. While the C-terminal region of MKSR-2 is not a recognised protein domain, it clearly has an important role in MKSR-2 organisation at the TZ. This observation is not surprising given that the C-terminus of MKSR-2 is highly conserved (**Figure 1A**).

### P74S and G155S mutations in MKSR-2 differentially disrupt cilia structure, positioning, and function

In *C. elegans*, the MKS and NPHP modules genetically interact to regulate cilium formation and function (Williams et al., 2008, 2011). For example, whereas cilia are mostly unaffected in single *mksr-2* and *nphp-4* null mutants, severe ciliogenesis defects and sensory behaviours (eg. osmotic avoidance) are observed in *mksr-2; nphp-4* double null mutants (Williams et al., 2008). Therefore, to investigate the phenotypic effects of the *mksr-2* patient alleles, we assessed sensory cilia in both wild-type/*nphp-4(+)* and *nphp-4(-)* genetic backgrounds.

First, we used a lipophilic dye (DiI/DiO) uptake assay to indirectly examine cilium integrity in 6 pairs of amphid (head) and both pairs of phasmid (tail) sensory neurons (Inglis et al., 2007; Starich et al., 1995). Defects in the structure of these environmentally exposed cilia strongly correlate with a dye filling (Dyf) defect (Inglis et al., 2007; Starich et al., 1995). Dyf phenotypes can be quantified by counting how many of the 4 phasmid cilia incorporate DiO. In the *nphp-4(+)* genetic background, none of the *mksr-2* alleles affect dye uptake (**Figure 3A, B**). However, in the *nphp-4(-)* background, the *mNG::mksr-2(P74S)* and *mNG::mksr-2(G155S)* alleles both cause very strong Dyf defects, which are almost as severe as that of the *mksr-2(Δ)* null allele (**Figure 3A, B**). The *mNG::mksr-2(P74S)* causes a slightly less severe phenotype compared to *mksr-2(G155S)* (p=0.006) (**Figure 3A, B**). The *mNG::mksr-2(+)* control allele causes a slight phasmid Dyf defect (avg 3.65 ± sd 0.7, p=0.006) indicating that the mNG tag negatively affects the *mksr-2* function to some extent in a minority of neurons.

**Figure 3.**
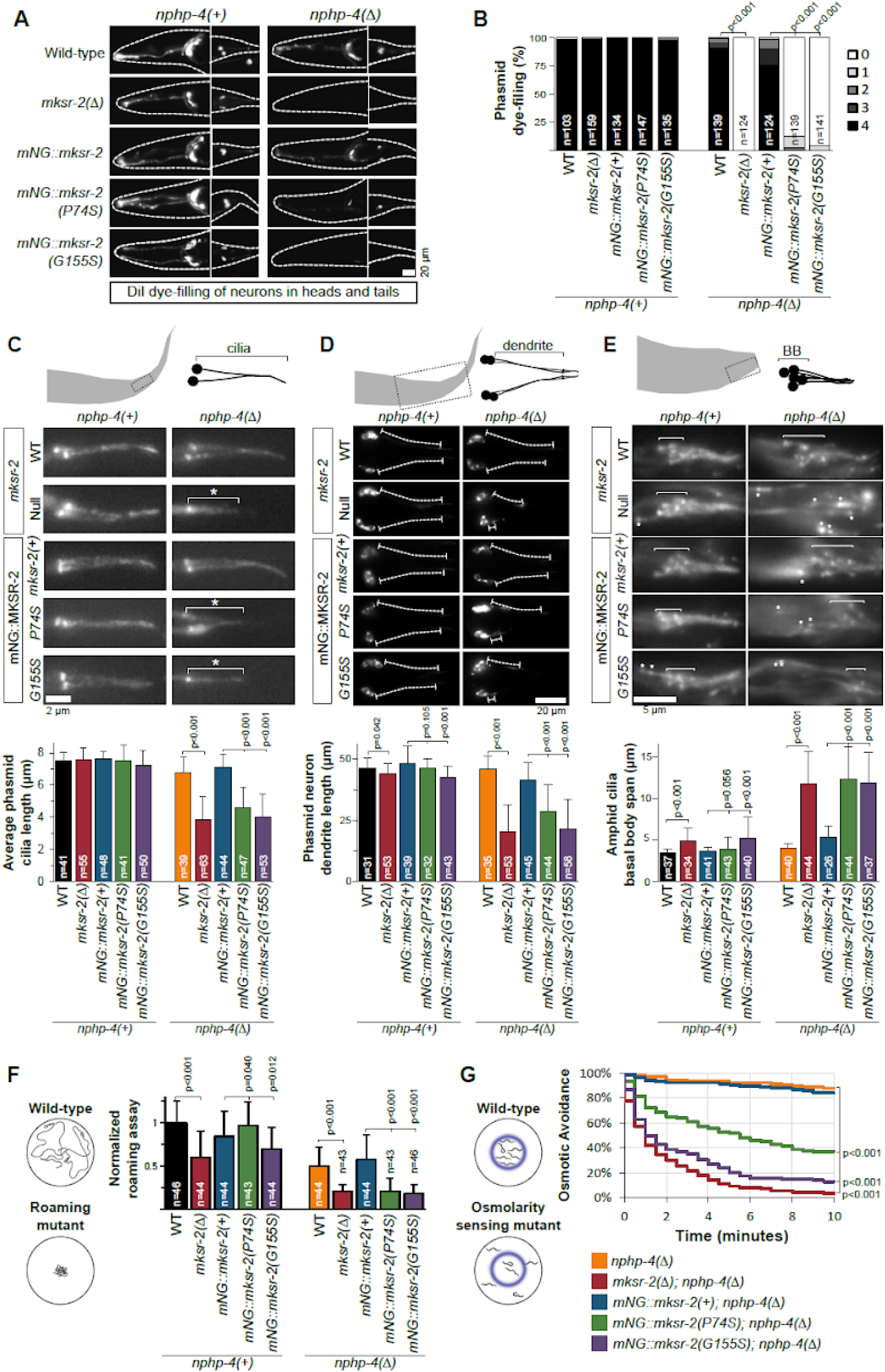
P74S and G155S mutations in MKSR-2 differentially disrupt cilium structure and function. **(A)** Representative images of DiI staining in head and tail ciliated sensory neurons of various *mksr-2* alleles in either *nphp-4(+)* or *nphp-4(-)* recessive genetic backgrounds. Dashed line indicates the worm body. Anterior is to the left. **(B)** Histogram showing the frequency of worms with DiI uptake in 0-4 phasmid neurons. Data is combined from three independent experiments. N; number of worms. p-values from unpaired t-tests. **(C, D)** Assessment of phasmid cilium length (C) and dendrite length (D) using an XBX-1::tdTomato reporter. Average cilia length calculated by measuring the XBX-1::tdTomato signal from the ciliary base to tip from a maximum projection. Dendrite length calculated by measuring the distance from the cell body to the base of the cilia from a maximum projection. Asterisk denotes short cilia. Dash lines denote the dendrite. Anterior is to the left. Scale bars; 2 μm (C) and 20 μm (D). Data in bar graphs represent mean and S.D. p-values from unpaired t-tests. N; number of cilia measured. **(E)** Assessment of amphid cilia clustering using the XBX-1::tdTomato reporter. Asterisks indicate amphid neuron BBs (and cilia) that are mis-positioned, proximal to the main cluster. Images are from a single plane. Anterior is to the right. Scale bars; 5 μm. Bar graphs quantify the distance spanned by amphid basal bodies from maximum projections. Data represent mean and S.D. p-values from unpaired t-tests. N; number of amphid pores. **(F)** Assessment of roaming behaviour. Data represent mean and S.D. p-values from unpaired t-tests. N; number of worms tested. Results normalised to wild-type. Data pooled from three independent experiments. **(G)** Assessment of osmotic avoidance behaviour in a 10 minute assay. n=18 from three independent experiments. p-values from unpaired t-tests.

Next, we employed an IFT-dynein light intermediate chain reporter (XBX-1::tdTomato) expressed in most or all sensory neurons to directly visualise cilia (Williams et al., 2011). First we measured the length of the phasmid cilia. As expected, in the *nphp-4(+)* background, the ~7.5 μm cilium length was not affected by any of the *mksr-2* alleles (**Figure 3C**). However, in the *nphp-4(-)* background, a 30-40% reduction in cilium length is associated with the *mNG::mksr-2(P74S)* (4.6 μm), *mNG::mksr-2(G155S)* (4.0 μm), and *mksr-2(Δ)* (3.9 μm) alleles, compared to the *mksr-2(+)* control (6.7 μm) (**Figure 3C**). When compared to *mksr-2(Δ)*, the cilium length phenotype associated with *mNG::mksr-2(G155S)* is equally severe (p=0.664), whilst that of *mNG::mksr-2* (P74S) is less severe (p=0.006). Using the XBX-1::tdTomato reporter, we also assessed cilium location, since loss of MKS module genes in an *nphp-4(-)* background disrupts the positioning of some cilia on account of abnormally short dendritic processes (Williams et al., 2008, 2011). For the 4 phasmid neurons, we measured dendrite length; for the 12 amphid neurons, we determined the spatial arrangement of their cilia by measuring the degree of basal body (BB) clustering. In the *nphp-4(+)* background, none of the *mksr-2* alleles dramatically affected cilium positioning, with the exception of a small increase in BB clustering caused by the *mksr-2(Δ)* and *mNG::mksr-2(G155S)* alleles (**Figure 3D, E**). In the *nphp-4(-)* background, and comparable to what is found for *mksr-2(Δ)*, the *mNG::mksr-2(G155S)* and *mNG::mksr-2(P74S)* alleles show a 40-50% reduction of phasmid dendrite length (**Figure 3D**) and amphid neuron BBs that are less tightly clustered (**Figure 3E**). Thus, both mutations disrupt cilium positioning.

Lastly, to determine if the *mksr-2* patient alleles affect cilia function we assessed two cilia-dependent sensory behaviours: foraging and osmotic avoidance. Wild-type *C. elegans* display a food foraging behaviour which can be quantified via a roaming assay (**Figure 3F**). In the *nphp-4(+)* background, the *mksr-2(Δ)* and *mNG::mksr-2(G155S)* alleles cause a reduction in roaming behaviour (**Figure 3F**). Note that a small roaming defect (p=0.008) is also found for the *mNG::mksr-2(+)* control allele (**Figure 3F**). In the *nphp-4(-)* background, the *mksr-2(Δ)*, *mNG::mksr-2(G155S)* and *mNG::mksr-2(P74S)* alleles all cause a similar strong roaming defect compared to controls (**Figure 3F**). In the osmotic avoidance assay, worms are assessed for their ability to avoid crossing a barrier (blue ring) of high osmolarity (**Figure 3G**). In the *nphp-4(-)* genetic background, the *mNG::mksr-2(G155S)* allele causes a strong reduction in osmotic sensing, almost as severe as that of the *mksr-2(Δ)* allele (Figure 3G). A reduced, albeit less severe, sensing defect is found for the *mNG::mksr-2(P74S)* allele (**Figure 3G**).

Together, these data show that the G155S and P74S mutations disrupt *mksr-2*’s role in cilium structure/function regulation. The findings also suggest that G155S is the more detrimental mutation, with phenotypes that are almost as severe as those of the null mutation.

### P74S and G155S mutations in MKSR-2 disrupt transition zone ultrastructure

We performed transmission electron microscopy to directly address the effects of the P74S and G155S *mksr-2* variants on TZ structure. As with general cilia structure phenotypes described above, defects in TZ structure and positioning are only observed in worms with null mutations in *nphp-4* and an MKS module gene (eg *mksr-1*), and not in most corresponding single mutants, with the exception of *cep-290* and *mks-5*. Thus, we analysed TZ ultrastructure in *mksr-2(P74S)*; *nphp-4(Δ)* and *mksr-2(G155S)*; *nphp-4(Δ)* double mutants. Specifically, we analysed serial thin sections of the bilateral amphid pore, each of which possesses 10 channel cilia extending from the distal dendrites of 8 amphid neurons (Doroquez et al., 2014; Perkins et al., 1986). These cilia consist of a degenerate basal body docked to the membrane of the periciliary membrane compartment (PCMC), followed by an ~0.8 μm long TZ, with Y-links connecting the 9 closely tethered outer doublet microtubules to the ciliary membrane (**Figure 4**). Following the TZ is the ~ 3 μm long middle segment, which consists of 9 doublet microtubules; the cilium ends with an ~ 3 μm long distal segment, comprising 9 singlet microtubules due to outer doublet B-tubule termination at the ends of the middle segment. Amphid channel cilia also contain varying numbers of inner singlet MTs.

**Figure 4.**
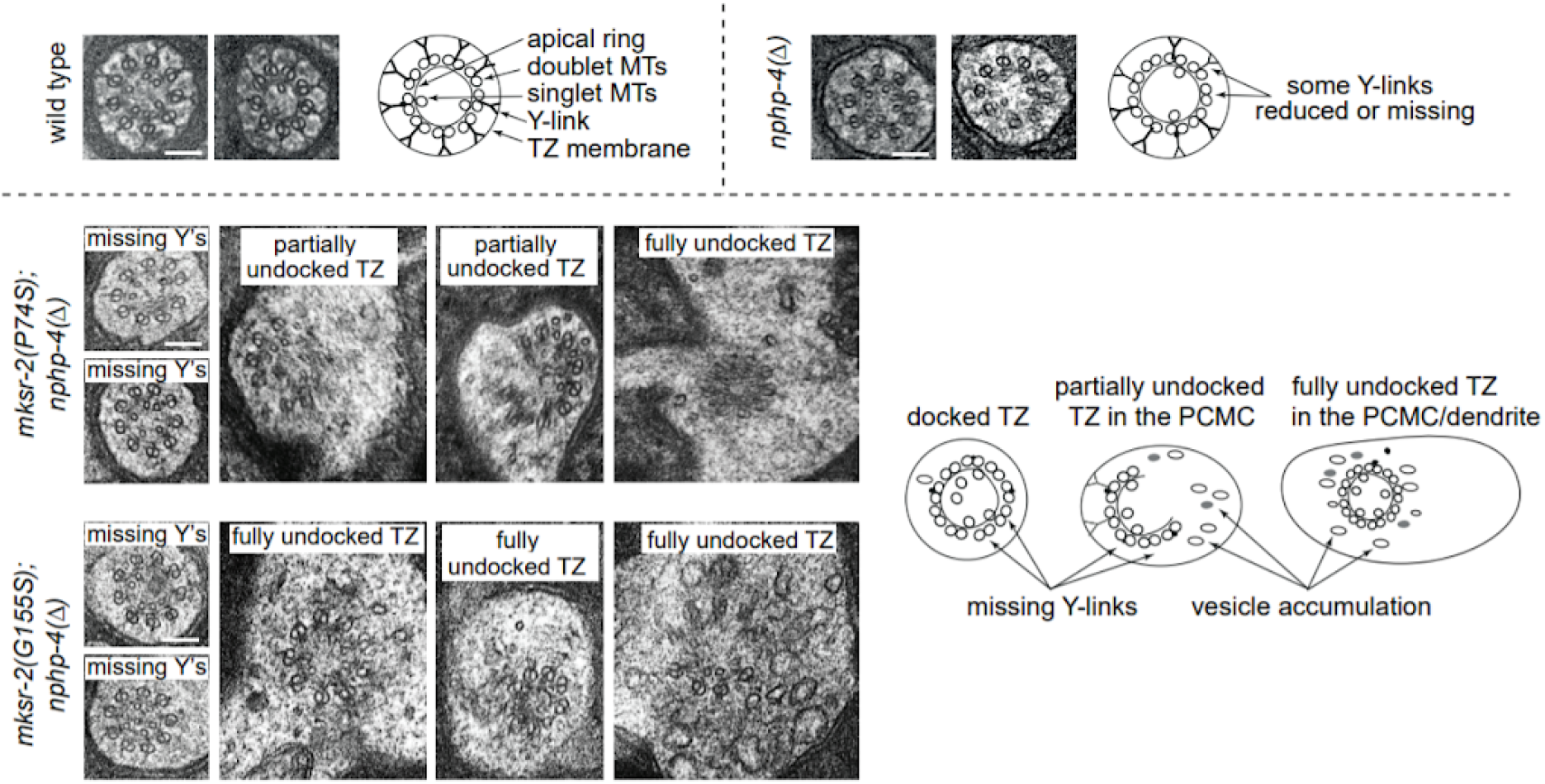
Ultrastructure of the transition zone is disrupted by P74S and G155S mutations in *mksr-2*. Transmission electron micrographs of amphid channel cilia transition zones in cross section (radial orientation) in *mksr-2(P74S)*; *nphp-4(Δ)* and *mksr-2(G155S)*; *nphp-4(Δ)* double mutants. Note that the wild-type and *nphp-4(Δ)* control images are adapted from (Lambacher et al., 2016). PCMC; periciliary membrane compartment. Y’s; Y-linkers. Scale bars; 100 nm (all images have the same scale).

The ultrastructure of *nphp-4*(*Δ*) (control) amphid channel cilia resembles that of wild type worms, except for a slight truncation of 1-2 cilia (**Figure S5; (Jauregui et al., 2008)**). *nphp-4*(*Δ*) worms also show a slight reduction in the density of some Y-linkers in the TZ, and in rare cases, the TZ is ectopically docked in more proximal positions of the PCMC or distal dendrite (**Figure 4; S5; (Lambacher et al., 2016)**). In contrast, both the *mksr-2(P74S)*; *nphp-4(Δ)* and *mksr-2(G155S)*; *nphp-4(Δ)* double mutants show major defects in cilia and TZ ultrastructure. Consistent with the light microscopy cilium structure phenotypes described above (**Figure 3E**), 4-7 amphid channel cilia are moderately or severely truncated; in some cases, the cilia are mispositioned in more proximal positions, extend at oblique angles to the other ciliary axonemes and are missing 1-3 MT doublets (**Figure S5**; data not shown). Y-links are not detectable in most TZs (**Figure 4**). In addition, many or most TZs do not extend from the periciliary membrane; instead, these ‘undocked’ TZ structures are present within the cytoplasm of more proximal positions of the PCMC and distal dendrite, which also often display a large accumulation of vesicles (**Figure 4**). Thus, the P74S and G155S mutations in *mksr-2* exert major defects on cilium and TZ ultrastructure. Indeed, the ultrastructure phenotype is similar to what we previously reported for null mutants of other core MKS module genes such as *mksr-1* (Williams et al., 2011).

### The G155S, but not P74S, mutation in MKSR-2 disrupts the ciliary gating of the *C. elegans* orthologue of Retinitis Pigmentosa 2 (RPI-2)

To assess how the G155S and P74S patient mutations affect the gating function of the ciliary TZ, we investigated their impact on cilium composition and the TZ membrane diffusion barrier. This was achieved using a fluorescent reporter of lipidated RPI-2 (orthologue of human Retinitis Pigmentosa 2), which is enriched at the periciliary membrane, but excluded from the ciliary membrane, ostensibly on account of a TZ membrane diffusion barrier (Williams et al., 2011). Unlike the cilium and TZ structure defects described above for double mutants of *nphp-4* and MKS module genes, defects in RPI-2 localisation are found in single mutants of these genes (Williams et al., 2011). Consistent with previous findings (Williams et al., 2011), we found that RPI-2::GFP expressed from an extrachromosomal array leaks into *mksr-2(Δ)* null cilia (**Figure 5A, B**). The RPI-2 reporter also leaks into the cilia of *mNG::mksr-2(G155S)* worms, albeit not to the same extent as in the null allele (**Figure 5A, B**). In contrast, the *mNG::mksr-2(P74S)* allele does not affect RPI-2 localisation when compared with the control (*mNG::mksr-2(+)*) (**Figure 5A, B**). Note that we are unable to distinguish between mNG::MKSR-2 and RPI-2::GFP fluorescence in this experiment. However, as mNG::MKSR-2 is never observed in the wild-type cilium distal to the TZ (**Figure 2A**), the green fluorescent signal observed in *mNG::mksr-2(G155S)* cilia is assumed to be RPI-2::GFP. Thus, the G155S mutation in MKSR-2 disrupts the ciliary gating of RPI-2, presumably by affecting the TZ membrane diffusion barrier.

**Figure 5.**
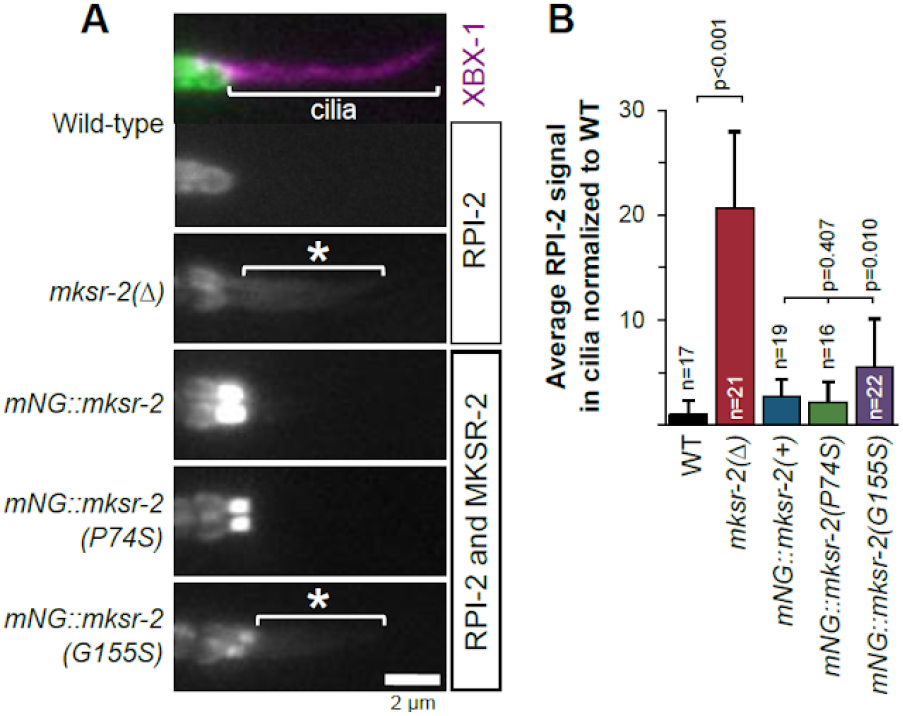
G155S, but not P74S, mutation in MKSR-2 causes the periciliary membrane protein RPI-2 to leak into cilia. **(A)** Representative images of phasmid cilia expressing the ciliary marker XBX-1::tdTomato and a reporter of the periciliary membrane protein, RPI-2::eGFP. Asterisk indicates RPI-2 signal in cilia. Note that XBX-1 signals are only shown in the top image. Anterior is to the left. Scale bar; 2 μm. **(B)** Quantification of RPI-2::GFP signal in phasmid cilia. Background fluorescence was subtracted and all samples are normalised to wild-type. Data represents mean with S.D. p-values from unpaired t-tests. N; number of phasmid cilia.

### TZ distribution of MKS-2 (core MKS module) is more severely affected than MKS-6 and JBTS-14 (peripheral MKS module) in worms with MKSR-2 patient mutations

MKSR-2 is part of the B9 complex (with MKS-1/MKS1 and B9D1/MKSR-1), which is a component of the core MKS module required for the TZ recruitment of other core and peripheral MKS module proteins including MKS-2/TMEM216 (core), JBST-14/TMEM237 (peripheral), and MKS-6/CC2D2A (peripheral); however, the core MKS module is not required for the TZ recruitment of the NPHP or CEP-290 modules, nor the master assembly factor MKS-5/RPGRIP1L (**Figure 6A**) (Huang et al., 2011; Jensen et al., 2015; Lambacher et al., 2016; C. Li et al., 2016; Williams et al., 2011). To assess whether the P74S and G155S mutations in MKSR-2 affect the recruitment of other TZ proteins we employed our untagged variant alleles: *mksr-2(oq137)*/*MKSR-2(P74S)* and *mksr-2(oq138)*/*MKSR-2(G155S)*. To examine the localisation of TZ module reporters expressed at endogenous levels, we used CRISPR-Cas9 editing to knock-in mNeonGreen at the *mks-5, nphp-4, cep-290, mks-2, jbts-14*, and *mks-6* loci. Using a dye-filling assay, it was found that the mNeonGreen tag did not appreciably disrupt the function of the associated genes, with the exception of a small effect on *mks-2* (**Figure S6A**).

**Figure 6.**
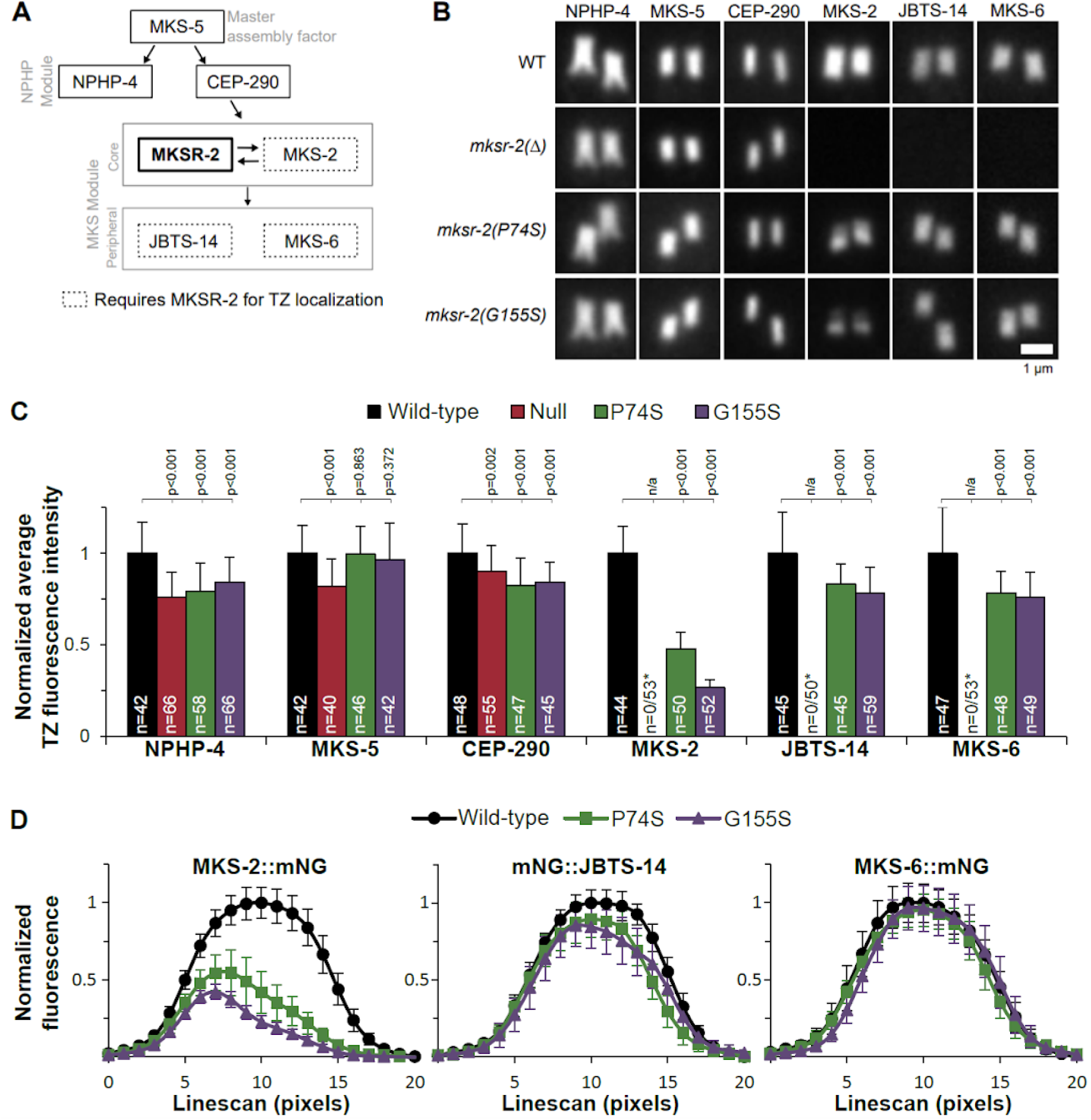
P74S and G155S mutations in MKSR-2 disrupt MKS module protein distribution at the transition zone to different extents. **(A)** Schematic showing the position and role of MKSR-2 within the hierarchical model of NPHP and MKS module assembly at the transition zone (TZ). MKSR-2 is a component of the core MKS module. Note: This is a simplified model with only 6 MKS/NPHP proteins shown. A more complete model can be found in Li *et al*., 2016. **(B)** Representative images of phasmid neuron TZs from worms expressing endogenous mNG::NPHP-4, mNG::MKS-5, CEP-290::mNG, MKS-2::mNG, mNG::JBTS-14, and MKS-6::mNG. Images taken from a single plane with both TZs in focus. Anterior is to the bottom. Scale bar; 1 μm. **(C)** Quantification of the total fluorescence intensity of mNeonGreen at the TZ of phasmid cilia in a single plane. Background fluorescence was subtracted and values normalised to wild-type. Data represent mean and S.D. p-values from unpaired t-tests. N; total number of phasmid pairs measured. Asterisk denotes that the fluorescence levels of MKS-2, JBTS-14, and MKS-6 in the *mksr-2* mutant background were not quantified because there is no TZ signal observed. **(D)** Average fluorescence intensity from line scans (20 pixels = 1.6 μm) of phasmid cilia showing the distribution of MKS-2, JBTS-14, and MKS-6 along the TZ length. Values normalised to wild-type. n=12 for each strain. Data represent mean +/- S.D.

First we examined endogenous reporter localisations in *mksr-2(Δ)* worms. Consistent with previous reports for core MKS module proteins, we found that the *mksr-2(Δ)* null mutant has lost TZ localisation of MKS-2, JBTS-14 and MKS-6, but retained TZ localisation of NPHP-4, CEP-290 and MKS-5 (**Figure 6B,C; S6D**). Two notable distinctions to reported data were observed. First, the mNG::MKS-2 reporter did not ectopically localise along the length of the *mksr-2(Δ)* mutant ciliary membrane as reported for an overexpressed MKS-2::GFP reporter (Huang et al., 2011). Second, the mNG::MKS-5 reporter localised as a continuous signal at the TZ of the *mksr-2(Δ)* mutant (**Figure S6B**), rather than an elongated signal with two clearly separated punctae as was found previously for an overexpressed MKS-5::tdTomato reporter in that mutant (Jensen et al., 2015). Indeed, a continuous mNG::MKS-5 signal is also found at the TZ region of *nphp-4, mks-2, and nphp-4; mks-2* mutants (**Figure S6C**), unlike what was observed in these mutants with MKS-5::tdTomato (Jensen et al., 2015). We conclude that these distinctions relate to the endogenous vs overexpressed nature of the reporters.

In contrast to the *mksr-2(Δ)* null allele, the endogenous mNG-tagged MKS-2, JBTS-14, and MKS-6 reporters are recruited to the ciliary TZs in *mksr-2(P74S)* and *mksr-2(G155S)* worms (**Figure 6B**). However, reporter fluorescence levels at the TZs are affected in these worms, albeit to different extents. Whereas mNG::JBST-14 and MKS-6::mNG signals are reduced by approximately 20%, the effect on MKS-2::mNG is more dramatic, showing a 50-75% reduction (most severe in G155S worms) (**Figure 6C**). Interestingly, in the *mksr-2* variants, the MKS-2::mNG signal is not evenly distributed at the TZ; instead, fluorescence clearly peaks at the proximal end (**Figure 6D**). Of note, the patient variants did not cause a dramatic proximal shift in the TZ localisation of JBTS-14 and MKS-6, although mNG::JBTS-14 shows a slight proximal shift in *mksr-2(G155S)* worms (**Figure 6D**). Together, these data show that the P74S and G155S mutations in MKSR-2 differentially disrupt the TZ localisations of core and peripheral MKS module proteins. Somewhat surprisingly, the TZ localisation of MKS-2 (core MKS module) is much more strongly affected than JBTS-14 and MKS-6 (peripheral MKS module).

### Using *C. elegans* genetic crosses to mimic ‘carrier’ and ‘pathogenic’ genotypes

Our analysis thus far has examined the MKSR-2 variants in a homozygous state. However, this does not accurately reflect the compound heterozygous genotype of the JBTS patient who inherited B9D2 variant alleles from carrier parents (**Figure 7A**) (R. Bachmann-Gagescu et al., 2015). Using standard genetic crossing techniques in *C. elegans*, we can investigate how these alleles behave as heterozygotes in the carrier and biallelic state (**Figure 7B**). A recessive phenotypic marker (*dpy-5*) was included in the crosses to verify all progeny analyzed were heterozygous. As described above, cilium structure and function assays were performed in an *nphp-4(-)* genetic background; controls are also heterozygous for the *dpy-5* allele. The complete genotypes of controls and heterozygotes are shown in **Figure S7**.

**Figure 7.**
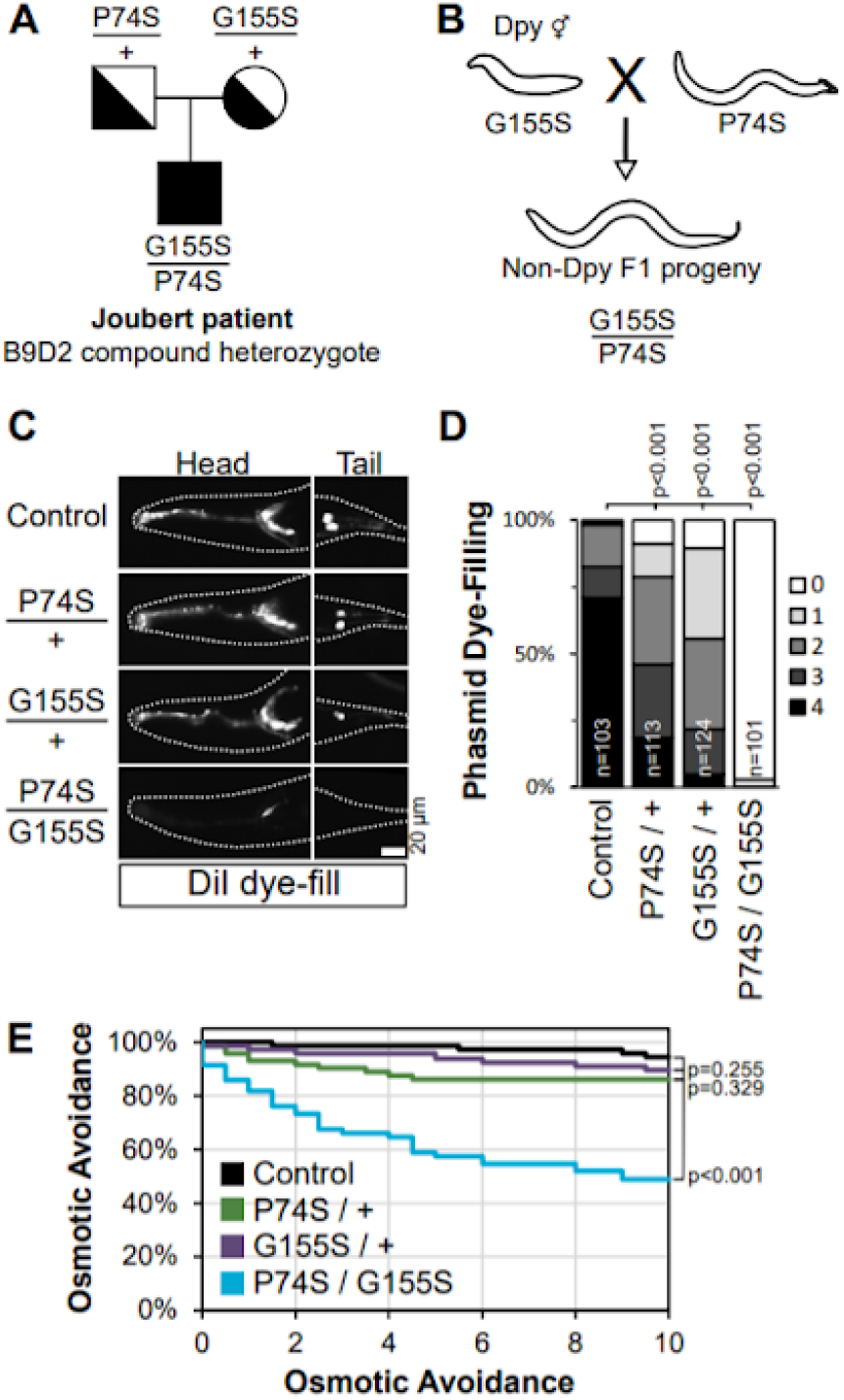
Phenotypic interpretation of *C. elegans* compound heterozygous G155S/P74S worms that mimics the biallelic genotype of a JBTS patient. **(A)** Pedigree of a Joubert syndrome biallelic patient described by Bachmann-Gagescu *et al*. (2015). Parents are heterozygous carriers of P74S or G155S. **(B)** Sample genetic cross illustrating how compound heterozygous F1 progeny are generated in *C. elegans*. A recessive *dpy-5* allele ensures analysis is only performed with outcrossed progeny. Full genotypes of genetic crosses used to generate heterozygous F1 worms are in Figure S7. **(C)** Representative dye uptake images in *mksr-2* heterozygotes that are homozygous for the *nphp-4* deletion allele. Anterior is to the left. Scale bar; 20 μm. **(D)** Quantification of dye uptake. Histogram shows the frequency of worms with dye uptake in 0-4 phasmid neurons. Data from three pooled independent experiments. N; number of worms. (E) Graphs of osmotic avoidance behaviour in 10 minute assays. p-values from unpaired t-tests, calculated for the 10 minute data points. n=12 from three independent experiments.

Using the standard lipophilic dye filling assay to assess phasmid cilium integrity we found that the heterozygous P74S/+ and G155S/+ carrier genotypes show a modest reduction in dye-filling (**Figure 7C, D**). In contrast, dye filling is completely abolished in compound heterozygous P74S/G155S worms (**Figure 7C, D**). Similarly, assessment of cilia function via the osmotic avoidance assay revealed grossly normal behaviours for heterozygous P74S/+ and G155S/+ worms, but highly defective behaviour for the compound heterozygous P74S/G155S animals (**Figure 7E**). These data are consistent with the recessive nature of the variants in Joubert Syndrome.

## DISCUSSION

### Modeling biallelic JBTS patient missense variants in *C. elegans* identifies differential effects for the P74S and G155S mutations on B9D2 gene function

In this study, we used CRISPR/Cas9 genome editing in the nematode *C. elegans* to model two pathogenic missense variants of *B9D2/mksr-2* associated with Joubert Syndrome. We found that in the homozygous state, both variants give rise to cilia and TZ defects that are found in a null allele of *mksr-2*. However, in most assays (cilia length, dye filling, dendrite length, and osmotic avoidance), the P74S variant phenotype was found to be milder than that of the G155S variant. A lipidated periciliary protein (RPI-2) is excluded from the cilia in the P74S but not G155S variant suggesting that the TZ diffusion barrier, at least to RPI-2, is disrupted only in the latter. Both variants showed a less severe overall phenotype compared to the null allele, although the latter was phenocopied by G155S in some assays. In line with these findings, super-resolution imaging revealed that the TZ localisation of the G155S variant protein is disorganised, whereas that of the P74S variant is more similar to wild-type. Together, these findings show that the G155S mutation has a greater effect on MKSR-2 gene function than the P74S mutation.

Our study also shows how *C. elegans* can be used to interpret more complex human genotypes such as the biallelic P74S/G155S genotype observed in a Joubert Syndrome patient. We found that P74S/G155S compound heterozygous worms exhibit more severe phenotypes than heterozygous carriers (P74S/+ and G155S/+) indicating that these alleles are recessive (**Figure 7C-E**), which is consistent with the previously described pedigree for these alleles (R. Bachmann-Gagescu et al., 2015). Additionally, we found that in heterozygous carriers the wild-type protein is more efficiently recruited to the TZ than the mutant protein (**Figure 2E**), thus providing a mechanism for the recessive nature of these variants. It is unclear why the P74S and G155S mutant proteins are not efficiently recruited to the TZ; one possibility is that the mutations could disrupt protein interactions in the B9 complex. Further experiments are required to assess this hypothesis.

We initially generated the P74S and G155S variants in an mNG::MKSR-2 strain for ease of assessing both cilium structure/function and localisation phenotypes. Throughout this study we observed that mNeonGreen on the N-terminus of the MKSR-2 protein sometimes exerted a mild defect on cilia structure and function (**Figure 3**). Therefore, for future studies of this nature, we recommend performing cilium structure/function assays on variants with the non-tagged endogenous gene and reserving knock-in tagged proteins solely for localisation studies.

Overall, our work demonstrates the utility of *C. elegans* for interpreting the effects of orthologous ciliopathy patient missense mutations on underlying gene function, at the level of cilium morphology, function and ultrastructure. Also, the modelling of more complex genotypes (eg. biallelic states) allows for an analysis that more accurately reflects the patient genotype.

### Hypomorphic patient alleles provide an opportunity to further refine the function of TZ functional modules

In *C. elegans*, most of our knowledge on TZ gene function has been derived using null alleles. For example, data from null alleles have established TZ localisation dependencies and assembly hierarchies for *C. elegans* NPHP and MKS module proteins (Bialas et al., 2009; Huang et al., 2011; Jensen et al., 2015; Lambacher et al., 2016; C. Li et al., 2016; Roberson et al., 2015; Schouteden et al., 2015; Williams et al., 2008, 2011; Winkelbauer et al., 2005; Yee et al., 2015). In the case of the MKS module, a core submodule that includes MKSR-2/B9D2 directs the TZ recruitment of a peripheral submodule (at least 6 proteins), but not vice versa. Therefore, it was surprising to find that in *mksr-2(P74S)* and *mksr-2(G155S)* worms the TZ level and distribution of a core MKS submodule component (MKS-2) was highly disrupted, whereas the TZ localisations of peripheral submodule proteins (JBTS-14/TMEM237, MKS-6/CC2D2A) were much less affected. This is in contrast to the *mksr-2* null allele, where MKS-2, MKS-6 and JBTS-14 all fail to assemble at the TZ (**Figure 6** and Huang et al., 2011; Williams et al., 2011). The lack of expected correlation for how peripheral and core MKS submodule components localise in *mksr-2(P74S)* and *mksr-2(G155S)* worms indicates that the relationship between these two submodules is more complex than previously thought. Our results demonstrate the benefits of using hypomorphic missense alleles to provide new biological insights that can not be elucidated with null alleles.

In addition, our data indicate an especially close functional relationship between MKSR-2 and MKS-2/TMEM216. Strikingly, we have found that MKS-2 redistribution to the proximal end of the TZ in *mksr-2(G155S)* worms is very similar to how the MKSR-2(G155S) protein itself is distributed at the TZ. Thus, TZ redistribution of MKSR-2 causes a comparable redistribution of MKS-2, suggesting a close spatial association of these proteins, perhaps as part of the same protein complex. In support of the latter, the mammalian B9 complex, consisting of MKS1, B9D1, and B9D2, has been shown to be in a larger complex with TMEM216, the mammalian orthologue of MKS-2 (Garcia-Gonzalo et al., 2011). However, it has also been found that the B9 complex interacts with CC2D2A (Garcia-Gonzalo et al., 2011; Gupta et al., 2015), the orthologue of MKS-6, which is in conflict with our observation of grossly normal MKS-6 localisations in the *mksr-2(G155S)* worms. Further research is required to determine how exactly these proteins are organised at the TZ.

### Endogenous and overexpressed reporters of MKS module proteins display several distinctions in their TZ localisation phenotype

In this study we examined the localisation of several fluorescent tagged TZ proteins, expressed at endogenous levels. Previous characterisation of TZ protein localisation in *C. elegans* employed transgenic proteins over-expressed from extrachromosomal arrays. Generally our results are consistent with previous observations, with a few notable exceptions.

Firstly, an overexpressed MKSR-2 reporter was previously reported to be localised to the TZ and at the distal end of the dendrite that includes the basal body and periciliary regions (Williams et al., 2008). This is in contrast to the endogenously expressed mNG::MKSR-2 protein, which is exclusively at the ciliary TZ, with no detectable pools in periciliary or dendritic regions. Secondly, it was previously reported that in null mutants of various MKS module genes (*mksr-1, mksr-2, mks-6, cep-290* and *tmem-231)*, an overexpressed MKS-2 reporter ectopically localises along the entire length of the cilium (Huang et al., 2011; C. Li et al., 2016; Roberson et al., 2015). In our experiments, we did not observe mislocalisation of the endogenous MKS-2::mNG reporter in the *mksr-2* null mutant (**Figure 6B, S5D**). Lastly, an overexpressed MKS-5::tdTomato reporter was previously found to form two discrete punctae of signal at either end (or possibly outside) the TZ in various MKS module gene mutants; however, in these same mutants, other MKS module proteins retain uniform signals along the TZ length (Jensen et al., 2015). This observation supported the conclusion that MKS-5 acts as an assembly factor for MKS module proteins, rather than a tethering scaffold, or structural anchor (Jensen et al., 2015). However, in our study here, the endogenous mNG::MKS-5 protein showed a TZ signal that was decreased in length and uniform in the *mksr-2* null mutant. This finding indicates that we cannot discount the notion that MKS-5 could act as a structural scaffold, and more research is required to answer this question.

Together, the above highlight important distinctions in the TZ localisation phenotype for overexpressed vs endogenously expressed reporters. Whilst the use of endogenous reporters is preferred for obvious reasons, distinct phenotypes arising from overexpressed reporters are still valuable for understanding TZ biology. For example, it would be interesting to further investigate how MKS-5 overexpression affects TZ ultrastructure: is the compartment elongated/continuous or is there a gap in the structure? Does excess MKS-5 recruit other TZ proteins to the proximal and distal regions of the TZ? Comparing and contrasting results obtained from overexpressed and endogenous level reporters can therefore provide novel insights into protein function and identify new avenues of investigation.

### *C. elegans* as a model to interrogate ciliopathy genotype-phenotype correlation and interpret variants of uncertain significance

Increasingly, whole genome or targeted gene panel sequencing is used to identify pathogenic variants in patients with ciliopathies (R. Bachmann-Gagescu et al., 2015; Wheway et al., 2019). If a pathogenic variant is determined to be the cause of a ciliopathy, this information can be used for genetic counselling, to adjust clinical care based on gene-specific complications, and even qualify patients for gene-specific clinical trials and treatments (Ruxandra Bachmann-Gagescu et al., 2020). Many sequence variants are termed “variants of unknown clinical significance” (VUS) because there is insufficient data for them to be classified as pathogenic or benign (M. M. Li et al., 2017). VUS alleles are not clinically useful and require further characterisation to be classified as pathogenic or benign. The data in this study shows that we can successfully model pathogenic ciliopathy mutations in *C. elegans*. Our work also shows that we can determine the precise effects of different missense mutations on the function of the orthologous gene by using quantitative assays of cilium structure and function, as well as knock-in fluorescent reporters for MKS module proteins expressed at the endogenous level. Since these assays are able to distinguish between the severity of the *mksr-2* alleles we tested, we anticipate they would serve as excellent tools to interpret ciliopathy VUS alleles. By modelling additional pathogenic alleles we will be able to determine the limits of our assays for interrogating patient allele pathogenicity based on *C. elegans* phenotypes. For example, do some known pathogenic alleles exhibit wild-type phenotypes in *C. elegans*? What is the rate of “false negatives” and “false positives” in the nematode assays? It will also be important to determine which assays fare better in stratifying the phenotypes of a wide spectrum of alleles at one locus, and whether data from multiple assays has greater predictive power for human genotype-phenotype correlation compared to a single assay.

Modelling ciliopathy patient variants in *C. elegans* has the potential to be a powerful diagnostic tool as there is a move towards personalised medicine for patients with rare genetic disorders. As shown here, patient variants can be efficiently generated in *C. elegans* using CRISPR-Cas9 technology, and rapidly identified using the simplified PCR-based allele detection strategy that we developed (**Figure 1S**). Indeed, with streamlined methodologies, it should be possible to generate hundreds of alleles in a relatively short time frame at relatively low cost and manpower compared to other multicellular systems. The workflow to generate and characterise ciliopathy associated variants described here can also be extended to other conserved cilia genes and ciliopathies. Of course, interpreting human missense variation in *C. elegans* is dependent on two key factors: (i) conservation of the mutated residue at the orthologous position in the nematode gene, and (ii) shared biology for the human and nematode genes. Ciliopathy genes are well served for these requirements; many of the genes show relatively high sequence identity to the worm counterpart and there is robust data showing comparable functions for human and *C. elegans* genes at the TZ. Given the shared biology, it might also be possible to humanise the nematode, by replacing the endogenous TZ gene with its human counterpart (McDiarmid et al., 2018). If the human orthologue retained functionality in the context of the worm TZ, this would allow for the testing of a much greater number of human variant mutations.

In conclusion, the efficiency of genetic engineering and relatively simple quantitative assays to characterise cilium structure and function in *C. elegans* makes it an excellent system for investigating genotype-phenotype correlation and interpreting VUS alleles in ciliopathies.

## MATERIALS AND METHODS

### *C. elegans* strains and maintenance

All *C. elegans* strains were maintained at 15°C or 20°C on NGM agar plates seeded with OP50 *E. coli* using standard worm culturing techniques (Brenner, 1974). PCR was used to confirm the genotypes of all strains. Young adults were synchronised by selecting L4 larvae and incubating at 20°C for 16-20 hours prior to imaging or phenotypic characterisation. A list of worm strains used in this study can be found in Table S2.

### Knock-in of mNeonGreen using CRISPR/Cas9

mNeonGreen (mNG) is licensed by Allele Biotechnology and Pharmaceuticals, Inc. (Shaner et al., 2013). A plasmid (dg353) containing *C. elegans* optimised mNeonGreen (Hostettler et al., 2017) was obtained from Dominique Glauser (University of Fribourg, Switzerland). High-fidelity PCR was used to add a 12aa flexible linker (GTGGGGSGGGGS) to the N- and C-terminus of mNG. PCR products were then cloned into pJET1.2 using the blunt end CloneJET PCR Cloning Kit (Thermo Scientific). Two rounds of high fidelity PCR was used to generate linear mNG repair templates with 35bp homology arms as previously described (Paix et al., 2015). All CRISPR protocols used Alt-R Cas9 nuclease 3NLS (IDT, #1074181), Alt-R tracrRNA (IDT, #1072533), and custom generated gene specific Alt-R crRNA from IDT. Gene specific crRNA sequences were chosen based on proximity to the desired edit and the Azimuth 2.0 score (Doench et al., 2016). All RNA was reconstituted with 5mM Tris pH 7.5 and stored at −80°C. ssODN repair templates were ordered as standard desalt DNA oligos from Sigma-Aldrich, reconstituted with 1M Tris pH 7.4, and stored at −20°C. A list of guide sequences and repair template sequences can be found in Table S1. All CRISPR injection mixes were prepared on ice, mixed gently, centrifuged at 13,000RPM for 2 minutes, and incubated at 37°C for 15 minutes prior to injection. N-terminal or C-terminal knock-ins of mNG were performed using a *dpy-10* co-CRISPR strategy (Paix et al., 2015). mNG knock-in injection mixes were as follows: 1 μl gene specific crRNA (0.3nmol/μl), 1 μl tracrRNA (0.425 nmol/μl), 0.2 μl *dpy-10* crRNA (0.6nmol/μl), 0.23 μl *dpy-10* ssODN (500 ng/μl), 5-6 μl mNG repair template (up to a maximum final concentration of 500 ng/μl), 1-2 μl 1M KCl, 0.38 μl HEPES (200mM, pH 7.4), 0.2 μl Cas9 (61μM), for a final volume of 10 μl. All Dpy and Rol F1 were screened in pools of 8 hermaphrodites. The knock-in efficiency of mNeonGreen varied from 0.1% to 2% with an average of 800 F1 screened per gene (range 376-2408). Sanger sequencing confirmed the accuracy of all knock-ins. Using this approach we generated *mks-2(oq101[MKS-2::mNG]), cep-290(oq103[CEP-290::mNG]), mksr-2(oq108[mNG::MKSR-2]), nphp-4(oq109[mNG::NPHP-4]), mks-5(oq112[mNG::MKS-5]), jbst-14(oq127[mNG::JBTS-14])*, and *mks-6(oq128[MKS-6::mNG])*.

### Engineering missense *mksr-2* variants with CRISPR/Cas9

The patient variants, *mksr-2(oq125[mNG::MKSR-2(P74S)])* and *mksr-2(oq126[mNG::MKSR-2(G155S)])*, were generated using ssODN repair templates, *unc-58* co-CRISPR (Arribere et al., 2014), and RFLP analysis to detect the variant (Paix et al., 2017). We modified the *unc-58* ssODN to include two additional silent mutations in the guide sequence (Table S1). *mksr-2(oq108[mNG::MKSR-2])* worms were used for the injections. crRNA and ssODN were reconstituted as described above. Injection mixes were prepared on ice as follows: 1 μl P74S crRNA (0.3 nmol/μl), 1 μl G155S crRNA (0.3 nmol/μl), 1 μl tracrRNA (0.425nmol/μl), 0.25 μl *unc-58* crRNA (1 nmol/μl), 0.25 μl *unc-58* ssODN (500 ng/μl), 0.5 μl P74S ssODN (1 μg/μl), 0.5 μl G155S ssODN (1 μg/μl), 2 μl 1M KCl, 0.38 μl HEPES (200 mM, pH 7.4), 0.2μl Cas9 (61 μM), and RNAse-free water up to 10 μl. The injection mix was incubated at 37°C for 15 minutes prior to injection. All Unc F1 were screened in pools of 1-4 hermaphrodites. The P74S and G155S variants introduced a restriction enzyme cut site in the *mksr-2* gene (BstUI and NheI, respectively). Using this restriction enzyme screening technique, the P74S and G155S variants were generated at an efficiency of 3.1% and 4.9%, respectively. The *mksr-2(oq137[P74S])* and *mksr-2(oq138[G155S]* strains were generated with the same approach and repair templates except wild-type N2 worms were used for the injections. These variants were detected in the F1 with a simplified PCR based approach using a primer that specifically binds to the engineered silent and missense mutations (Figure S1). Both variants were generated with an efficiency of 12.5%. Sanger sequencing confirmed the accuracy of all CRISPR generated alleles and each strain was outcrossed twice prior to analysis.

### Wide-field epifluorescence live imaging and quantification

Worms were immobilised in 40mM tetramisole (Sigma) on 4% or 10% agarose pads. All images were acquired on an upright Leica DM5000B epifluorescence microscope. Images were collected with 100x (1.40 NA) or 63x (1.3 NA) oil-immersion objectives and captured with an Andor iXon+ camera run by Andor software. All image analysis and preparation was performed with FIJI/ImageJ (NIH). The TZ length of the mNG::MKSR-2 or mNG::MKS-5 signal length in phasmid neurons was calculated using a ‘full width at half max’ approach from line scans at the center of the TZ from single focal planes. The fluorescence levels of mNeonGreen tagged proteins at the TZ were quantified from a single focal plane where both TZs were in focus. The integrated signal intensity in a 40×40 pixel box around the TZ pair was measured and then increased by 1 pixel in each direction. The integrated signal intensity of this 42×42 pixel box was used to calculate and subtract background fluorescence. Phasmid cilia and dendrite lengths were measured from maximum projections of XBX-1::tdTomato using the segmented line feature of ImageJ. The span of the amphid BBs were measured from maximum projections with a box that was drawn around the BBs. To measure RPI-2::GFP levels in the cilia a segmented line was drawn along the cilia in the XBX-1::tdTomato channel from the base to beyond the cilia tip. RPI-2::GFP levels 4 μm from the base were used in calculations. The background was calculated by averaging the fluorescence intensity values at the end of the line beyond the cilia tip.

### Confocal super-resolution microscopy

Confocal based super-resolution microscopy was achieved with a reduced pin hole size and deconvolution (Lam et al., 2017). Super-resolution imaging was performed on an Olympus FV3000 confocal laser scanning microscope with a 60x (NA 1.40) oil-immersion objective and high-sensitivity spectral detector using FluoView Olympus software. The Airy disk pinhole size was set to 0.4 (81 μm) and 15x optical zoom was utilised. The pixel size was 26.7 nm with an X-Y optical resolution of 148.6 nm. Z-stacks were acquired with 0.2 μm steps. All images were deconvoled with Olympus cellSens software using default values. Imaris 7.6.5 was used to generate three-dimensional models. To account for the lower z-plane resolution a step size of 0.1 μm was used to create the 3D models.

### Assays to characterise cilia defects: dye filling, roaming, and osmotic avoidance

All phenotypic characterisation of cilia defects was performed as previously described (Sanders et al., 2015) with young adult hermaphrodites. Dye filling assays were performed with worms that were incubated in a 1 in 200 dilution of DiI or DiO (Invitrogen) in M9 for 60 minutes then allowed to recover for 30 minutes on seeded NGM plates. Worms were then mounted on 4% agarose pads with 40 mM tetramisole (Sigma) to assess dye uptake with a wide-field epifluorescence microscope. DiI (red) was used for image acquisition and DiO (green) was used for quantification. The assay was repeated at least 3 times with 30-50 worms per trial. Roaming assays were performed with a single worm that was placed on a fully seeded NGM plate for 20 hours at 20°C. The worm was removed and the plate was placed on a 5 x 5 mm grid and the number of squares that the worm entered was counted. For each strain tested, the assay was repeated at least 3 times with 15-20 worms per trial. Dye-filling and roaming assays were performed blindly by randomizing the strains before the start of the experiment. For the osmotic avoidance assay worms were placed on an unseeded NGM plate for 2 minutes prior to the assay. 6 worms were placed in a high osmolarity ring (8M glycerol with Bromophenol Blue) and observed for 10 minutes. The time when any worms escaped the barrier was noted and the worm was removed from the assay. Three independent trials were performed for each osmotic avoidance assay.

### Transmission electron microscopy

Young adult hermaphrodites were fixed, embedded, sectioned and imaged as previously described (Sanders et al., 2015). Briefly, worms were fixed in 2.5% glutaraldehyde in Sørensen’s phosphate buffer (0.1M, pH7.4) for 48 hours at 4°C. Samples were post-fixed in 1% osmium tetroxide for 1 hour and dehydrated through an increasing ethanol gradient. Ethanol was substituted for propylene oxide prior to EPON resin embedding overnight. Serial, ultra-thin 90 nm sections were cut using a Leica EM UC6 Ultramicrotome, collected on copper grids and subjected to double contrasting using 2% uranyl acetate for 20 minutes followed by 3% lead citrate for 5 minutes. Imaging was performed on a Tecnai 12 (FEI software) under acceleration voltage of 120 kV.

### Source Data - All Figures

Supplemental spreadsheet containing numerical data for all graphs with statistical analysis including Figures 2B, 2C, 2G, 3B, 3C, 3D, 3E, 3F, 3G, 5B, 6C, 7D, 7E, S3B, S6A, and S6B.

## ACKNOWLEDGEMENTS

This work was funded by Science Foundation Ireland (SFI) in partnership with the Biotechnology and Biological Sciences Research Council (BBSRC) under grant number 16/BBSRC/3394. We thank the UCD Conway Institute imaging facility (Dimitri Scholz, Tiina O’Neill, Niamh Stephens) for imaging assistance and Craig McCauley for help writing scripts to streamline analysis in Image J. We thank the Caenorhabditis Genetics Center (CGC) and National Bioresource Project (Tokyo Women’s Medical College, Tokyo, Japan). The CGC is supported by the National Institutes of Health - Office of Research Infrastructure Programs (P40 OD010440).

## REFERENCES

Anvarian, Z., Mykytyn, K., Mukhopadhyay, S., Pedersen, L. B., & Christensen, S. T. (2019). Cellular signalling by primary cilia in development, organ function and disease. Nature Reviews. Nephrology, 15(4), 199–219.

Arribere, J. A., Bell, R. T., Fu, B. X. H., Artiles, K. L., Hartman, P. S., & Fire, A. Z. (2014). Efficient marker-free recovery of custom genetic modifications with CRISPR/Cas9 in Caenorhabditis elegans. Genetics, 198(3), 837–846.

Bachmann-Gagescu, R., Dempsey, J. C., Bulgheroni, S., Chen, M. L., D’Arrigo, S., Glass, I. A., Heller, T., Héon, E., Hildebrandt, F., Joshi, N., Knutzen, D., Kroes, H. Y., Mack, S. H., Nuovo, S., Parisi, M. A., Snow, J., Summers, A. C., Symons, J. M., Zein, W. M., … Doherty, D. (2020). Healthcare recommendations for Joubert syndrome. American Journal of Medical Genetics. Part A, 182(1), 229–249.

Bachmann-Gagescu, R., Dempsey, J. C., Phelps, I. G., O’Roak, B. J., Knutzen, D. M., Rue, T. C., Ishak, G. E., Isabella, C. R., Gorden, N., Adkins, J., Boyle, E. A., de Lacy, N., O’Day, D., Alswaid, A., Ramadevi A, R., Lingappa, L., Lourenço, C., Martorell, L., Garcia-Cazorla, À., … Doherty, D. (2015). Joubert syndrome: a model for untangling recessive disorders with extreme genetic heterogeneity. Journal of Medical Genetics, 52(8), 514–522.

Bader, I., Decker, E., Mayr, J. A., Lunzer, V., Koch, J., Boltshauser, E., Sperl, W., Pietsch, P., Ertl-Wagner, B., Bolz, H., Bergmann, C., & Rittinger, O. (2016). MKS1 mutations cause Joubert syndrome with agenesis of the corpus callosum. European Journal of Medical Genetics, 59(8), 386–391.

Basiri, M. L., Ha, A., Chadha, A., Clark, N. M., Polyanovsky, A., Cook, B., & Avidor-Reiss, T. (2014). A migrating ciliary gate compartmentalizes the site of axoneme assembly in Drosophila spermatids. Current Biology: CB, 24 (22), 2622–2631.

Bialas, N. J., Inglis, P. N., Li, C., Robinson, J. F., Parker, J. D. K., Healey, M. P., Davis, E. E., Inglis, C. D., Toivonen, T., Cottell, D. C., Blacque, O. E., Quarmby, L. M., Katsanis, N., & Leroux, M. R. (2009). Functional interactions between the ciliopathy-associated Meckel syndrome 1 (MKS1) protein and two novel MKS1-related (MKSR) proteins. Journal of Cell Science, 122 (Pt 5), 611–624.

Blacque, O. E., & Sanders, A. A. W. M. (2014). Compartments within a compartment: what C. elegans can tell us about ciliary subdomain composition, biogenesis, function, and disease. Organogenesis, 10(1), 126–137.

Boldt, K., van Reeuwijk, J., Lu, Q., Koutroumpas, K., Nguyen, T.-M. T., Texier, Y., van Beersum, S. E. C., Horn, N., Willer, J. R., Mans, D. A., Dougherty, G., Lamers, I. J. C., Coene, K. L. M., Arts, H. H., Betts, M. J., Beyer, T., Bolat, E., Gloeckner, C. J., Haidari, K., … UK10K Rare Diseases Group. (2016). An organelle-specific protein landscape identifies novel diseases and molecular mechanisms. Nature Communications, 7, 11491.

Brenner, S. (1974). The genetics of Caenorhabditis elegans. Genetics, 77(1), 71–94.

Cevik, S., Sanders, A. A. W. M., Van Wijk, E., Boldt, K., Clarke, L., van Reeuwijk, J., Hori, Y., Horn, N., Hetterschijt, L., Wdowicz, A., Mullins, A., Kida, K., Kaplan, O. I., van Beersum, S. E. C., Man Wu, K., Letteboer, S. J. F., Mans, D. A., Katada, T., Kontani, K., … Blacque, O. E. (2013). Active transport and diffusion barriers restrict Joubert Syndrome-associated ARL13B/ARL-13 to an Inv-like ciliary membrane subdomain. PLoS Genetics, 9(12), e1003977.

Chih, B., Liu, P., Chinn, Y., Chalouni, C., Komuves, L. G., Hass, P. E., Sandoval, W., & Peterson, A. S. (2011). A ciliopathy complex at the transition zone protects the cilia as a privileged membrane domain. Nature Cell Biology, 14(1), 61–72.

Cui, C., Chatterjee, B., Francis, D., Yu, Q., SanAgustin, J. T., Francis, R., Tansey, T., Henry, C., Wang, B., Lemley, B., Pazour, G. J., & Lo, C. W. (2011). Disruption of Mks1 localization to the mother centriole causes cilia defects and developmental malformations in Meckel-Gruber syndrome. Disease Models & Mechanisms, 4(1), 43–56.

Doench, J. G., Fusi, N., Sullender, M., Hegde, M., Vaimberg, E. W., Donovan, K. F., Smith, I., Tothova, Z., Wilen, C., Orchard, R., Virgin, H. W., Listgarten, J., & Root, D. E. (2016). Optimized sgRNA design to maximize activity and minimize off-target effects of CRISPR-Cas9. Nature Biotechnology, 34(2), 184–191.

Doroquez, D. B., Berciu, C., Anderson, J. R., Sengupta, P., & Nicastro, D. (2014). A high-resolution morphological and ultrastructural map of anterior sensory cilia and glia in Caenorhabditis elegans. eLife, 3, e01948.

Dowdle, W. E., Robinson, J. F., Kneist, A., Sirerol-Piquer, M. S., Frints, S. G. M., Corbit, K. C., Zaghloul, N. A., van Lijnschoten, G., Mulders, L., Verver, D. E., Zerres, K., Reed, R. R., Attié-Bitach, T., Johnson, C. A., García-Verdugo, J. M., Katsanis, N., Bergmann, C., & Reiter, J. F. (2011). Disruption of a ciliary B9 protein complex causes Meckel syndrome. American Journal of Human Genetics, 89(1), 94–110.

El-Brolosy, M. A., Kontarakis, Z., Rossi, A., Kuenne, C., Günther, S., Fukuda, N., Kikhi, K., Boezio, G. L. M., Takacs, C. M., Lai, S.-L., Fukuda, R., Gerri, C., Giraldez, A. J., & Stainier, D. Y. R. (2019). Genetic compensation triggered by mutant mRNA degradation. Nature, 568(7751), 193–197.

Ganner, A., & Neumann-Haefelin, E. (2017). Genetic kidney diseases: Caenorhabditis elegans as model system. Cell and Tissue Research, 369(1), 105–118.

Garcia-Gonzalo, F. R., Corbit, K. C., Sirerol-Piquer, M. S., Ramaswami, G., Otto, E. A., Noriega, T. R., Seol, A. D., Robinson, J. F., Bennett, C. L., Josifova, D. J., García-Verdugo, J. M., Katsanis, N., Hildebrandt, F., & Reiter, J. F. (2011). A transition zone complex regulates mammalian ciliogenesis and ciliary membrane composition. Nature Genetics, 43 (8), 776–784.

Garcia-Gonzalo, F. R., & Reiter, J. F. (2017). Open Sesame: How Transition Fibers and the Transition Zone Control Ciliary Composition. Cold Spring Harbor Perspectives in Biology, 9(2). https://doi.org/10.1101/cshperspect.a028134

Gupta, G. D., Coyaud, É., Gonçalves, J., Mojarad, B. A., Liu, Y., Wu, Q., Gheiratmand, L., Comartin, D., Tkach, J. M., Cheung, S. W. T., Bashkurov, M., Hasegan, M., Knight, J. D., Lin, Z.-Y., Schueler, M., Hildebrandt, F., Moffat, J., Gingras, A.-C., Raught, B., & Pelletier, L. (2015). A Dynamic Protein Interaction Landscape of the Human Centrosome-Cilium Interface. Cell, 163(6), 1484–1499.

Hostettler, L., Grundy, L., Käser-Pébernard, S., Wicky, C., Schafer, W. R., & Glauser, D. A. (2017). The Bright Fluorescent Protein mNeonGreen Facilitates Protein Expression Analysis In Vivo. G3, 7(2), 607–615.

Huang, L., Szymanska, K., Jensen, V. L., Janecke, A. R., Innes, A. M., Davis, E. E., Frosk, P., Li, C., Willer, J. R., Chodirker, B. N., Greenberg, C. R., McLeod, D. R., Bernier, F. P., Chudley, A. E., Müller, T., Shboul, M., Logan, C. V., Loucks, C. M., Beaulieu, C. L., … Boycott, K. M. (2011). TMEM237 is mutated in individuals with a Joubert syndrome related disorder and expands the role of the TMEM family at the ciliary transition zone. American Journal of Human Genetics, 89(6), 713–730.

Inglis, P. N., Ou, G., Leroux, M. R., & Scholey, J. M. (2007). The sensory cilia of Caenorhabditis elegans. WormBook: The Online Review of C. Elegans Biology, 1–22.

Jauregui, A. R., Nguyen, K. C. Q., Hall, D. H., & Barr, M. M. (2008). The Caenorhabditis elegans nephrocystins act as global modifiers of cilium structure. The Journal of Cell Biology, 180 (5), 973–988.

Jensen, V. L., & Leroux, M. R. (2017). Gates for soluble and membrane proteins, and two trafficking systems (IFT and LIFT), establish a dynamic ciliary signaling compartment. Current Opinion in Cell Biology, 47, 83–91.

Jensen, V. L., Li, C., Bowie, R. V., Clarke, L., Mohan, S., Blacque, O. E., & Leroux, M. R. (2015). Formation of the transition zone by Mks5/Rpgrip1L establishes a ciliary zone of exclusion (CIZE) that compartmentalises ciliary signalling proteins and controls PIP2 ciliary abundance. The EMBO Journal, 34(20) 2537–2556.

Kim, W., Underwood, R. S., Greenwald, I., & Shaye, D. D. (2018). OrthoList 2: A New Comparative Genomic Analysis of Human and Caenorhabditis elegans Genes. Genetics, 210(2), 445–461.

Lambacher, N. J., Bruel, A.-L., van Dam, T. J. P., Szymańska, K., Slaats, G. G., Kuhns, S., McManus, G. J., Kennedy, J. E., Gaff, K., Wu, K. M., van der Lee, R., Burglen, L., Doummar, D., Rivière, J.-B., Faivre, L., Attié-Bitach, T., Saunier, S., Curd, A., Peckham, M., … Blacque, O. E. (2016). TMEM107 recruits ciliopathy proteins to subdomains of the ciliary transition zone and causes Joubert syndrome. Nature Cell Biology, 18(1), 122–131.

Lam, F., Cladière, D., Guillaume, C., Wassmann, K., & Bolte, S. (2017). Super-resolution for everybody: An image processing workflow to obtain high-resolution images with a standard confocal microscope. Methods, 115, 17–27.

Lemmon, M. A. (2008). Membrane recognition by phospholipid-binding domains. Nature Reviews. Molecular Cell Biology, 9 (2), 99–111.

Lewis, W. R., Bales, K. L., Revell, D. Z., Croyle, M. J., Engle, S. E., Song, C. J., Malarkey, E. B., Uytingco, C. R., Shan, D., Antonellis, P. J., Nagy, T. R., Kesterson, R. A., Mrug, M. M., Martens, J. R., Berbari, N. F., Gross, A. K., & Yoder, B. K. (2019). Mks6 mutations reveal tissue- and cell type-specific roles for the cilia transition zone. FASEB Journal: Official Publication of the Federation of American Societies for Experimental Biology, 33(1), 1440–1455.

Li, C., Jensen, V. L., Park, K., Kennedy, J., Garcia-Gonzalo, F. R., Romani, M., De Mori, R., Bruel, A.-L., Gaillard, D., Doray, B., Lopez, E., Rivière, J.-B., Faivre, L., Thauvin-Robinet, C., Reiter, J. F., Blacque, O. E., Valente, E. M., & Leroux, M. R. (2016). MKS5 and CEP290 Dependent Assembly Pathway of the Ciliary Transition Zone. PLoS Biology, 14(3), e1002416.

Li, M. M., Datto, M., Duncavage, E. J., Kulkarni, S., Lindeman, N. I., Roy, S., Tsimberidou, A. M., Vnencak-Jones, C. L., Wolff, D. J., Younes, A., & Nikiforova, M. N. (2017). Standards and Guidelines for the Interpretation and Reporting of Sequence Variants in Cancer: A Joint Consensus Recommendation of the Association for Molecular Pathology, American Society of Clinical Oncology, and College of American Pathologists. The Journal of Molecular Diagnostics: JMD, 19(1), 4–23.

Li, S., Armstrong, C. M., Bertin, N., Ge, H., Milstein, S., Boxem, M., Vidalain, P.-O., Han, J.-D. J., Chesneau, A., Hao, T., Goldberg, D. S., Li, N., Martinez, M., Rual, J.-F., Lamesch, P., Xu, L., Tewari, M., Wong, S. L., Zhang, L. V., … Vidal, M. (2004). A Map of the Interactome Network of the Metazoan C. elegans. Science, 303(5657), 540–543.

McDiarmid, T. A., Au, V., Loewen, A. D., Liang, J., Mizumoto, K., Moerman, D. G., & Rankin, C. H. (2018). CRISPR-Cas9 human gene replacement and phenomic characterization in Caenorhabditis elegans to understand the functional conservation of human genes and decipher variants of uncertain significance. Disease Models & Mechanisms, 11(12). https://doi.org/10.1242/dmm.036517

Mok, C. A., & Héon, E. (2012). Caenorhabditis elegans as a model organism for ciliopathies and related forms of photoreceptor degeneration. Advances in Experimental Medicine and Biology, 723, 533–538.

Molinari, E., & Sayer, J. A. (2020). Disease Modeling To Understand the Pathomechanisms of Human Genetic Kidney Disorders. Clinical Journal of the American Society of Nephrology: CJASN. https://doi.org/10.2215/CJN.08890719

Nachury, M. V., & Mick, D. U. (2019). Establishing and regulating the composition of cilia for signal transduction. Nature Reviews. Molecular Cell Biology, 20(7), 389–405.

Paix, A., Folkmann, A., Rasoloson, D., & Seydoux, G. (2015). High Efficiency, Homology-Directed Genome Editing in Caenorhabditis elegans Using CRISPR-Cas9 Ribonucleoprotein Complexes. Genetics, 201(1), 47–54.

Paix, A., Folkmann, A., & Seydoux, G. (2017). Precision genome editing using CRISPR-Cas9 and linear repair templates in C. elegans. Methods, 121-122, 86–93.

Perkins, L. A., Hedgecock, E. M., Thomson, J. N., & Culotti, J. G. (1986). Mutant sensory cilia in the nematode Caenorhabditis elegans. Developmental Biology, 117(2), 456–487.

Ponsard, C., Skowron-Zwarg, M., Seltzer, V., Perret, E., Gallinger, J., Fisch, C., Dupuis-Williams, P., Caruso, N., Middendorp, S., & Tournier, F. (2007). Identification of ICIS-1, a new protein involved in cilia stability. Frontiers in Bioscience: A Journal and Virtual Library, 12, 1661–1669.

Reiter, J. F., Blacque, O. E., & Leroux, M. R. (2012). The base of the cilium: roles for transition fibres and the transition zone in ciliary formation, maintenance and compartmentalization. EMBO Reports, 13(7), 608–618.

Reiter, J. F., & Leroux, M. R. (2017). Genes and molecular pathways underpinning ciliopathies. Nature Reviews. Molecular Cell Biology, 18(9), 533–547.

Roberson, E. C., Dowdle, W. E., Ozanturk, A., Garcia-Gonzalo, F. R., Li, C., Halbritter, J., Elkhartoufi, N., Porath, J. D., Cope, H., Ashley-Koch, A., Gregory, S., Thomas, S., Sayer, J. A., Saunier, S., Otto, E. A., Katsanis, N., Davis, E. E., Attié-Bitach, T., Hildebrandt, F., … Reiter, J. F. (2015). TMEM231, mutated in orofaciodigital and Meckel syndromes, organizes the ciliary transition zone. The Journal of Cell Biology, 209 (1), 129–142.

Romani, M., Micalizzi, A., Kraoua, I., Dotti, M. T., Cavallin, M., Sztriha, L., Ruta, R., Mancini, F., Mazza, T., Castellana, S., Hanene, B., Carluccio, M. A., Darra, F., Máté, A., Zimmermann, A., Gouider-Khouja, N., & Valente, E. M. (2014). Mutations in B9D1 and MKS1 cause mild Joubert syndrome: expanding the genetic overlap with the lethal ciliopathy Meckel syndrome. Orphanet Journal of Rare Diseases, 9, 72.

Sanders, A. A. W. M., Anna A W, Kennedy, J., & Blacque, O. E. (2015). Image analysis of Caenorhabditis elegans ciliary transition zone structure, ultrastructure, molecular composition, and function. In Methods in Cilia & Flagella (pp. 323–347). https://doi.org/10.1016/bs.mcb.2015.01.010

Satir, P., Pedersen, L. B., & Christensen, S. T. (2010). The primary cilium at a glance. In Journal of Cell Science (Vol. 123, Issue 4, pp. 499–503). https://doi.org/10.1242/jcs.050377

Schouteden, C., Serwas, D., Palfy, M., & Dammermann, A. (2015). The ciliary transition zone functions in cell adhesion but is dispensable for axoneme assembly in C. elegans. The Journal of Cell Biology, 210(1), 35–44.

Shaner, N. C., Lambert, G. G., Chammas, A., Ni, Y., Cranfill, P. J., Baird, M. A., Sell, B. R., Allen, J. R., Day, R. N., Israelsson, M., Davidson, M. W., & Wang, J. (2013). A bright monomeric green fluorescent protein derived from Branchiostoma lanceolatum. Nature Methods, 10 (5), 407–409.

Shimada, H., Lu, Q., Insinna-Kettenhofen, C., Nagashima, K., English, M. A., Semler, E. M., Mahgerefteh, J., Cideciyan, A. V., Li, T., Brooks, B. P., Gunay-Aygun, M., Jacobson, S. G., Cogliati, T., Westlake, C. J., & Swaroop, A. (2017). In Vitro Modeling Using Ciliopathy-Patient-Derived Cells Reveals Distinct Cilia Dysfunctions Caused by CEP290 Mutations. Cell Reports, 20(2), 384–396.

Shi, X., Garcia, G., 3rd, Van De Weghe, J. C., McGorty, R., Pazour, G. J., Doherty, D., Huang, B., & Reiter, J. F. (2017). Super-resolution microscopy reveals that disruption of ciliary transition-zone architecture causes Joubert syndrome. Nature Cell Biology, 19(10), 1178–1188.

Simonis, N., Rual, J.-F., Carvunis, A.-R., Tasan, M., Lemmens, I., Hirozane-Kishikawa, T., Hao, T., Sahalie, J. M., Venkatesan, K., Gebreab, F., Cevik, S., Klitgord, N., Fan, C., Braun, P., Li, N., Ayivi-Guedehoussou, N., Dann, E., Bertin, N., Szeto, D., … Vidal, M. (2009). Empirically controlled mapping of the Caenorhabditis elegans protein-protein interactome network. Nature Methods, 6(1), 47–54.

Slaats, G. G., Isabella, C. R., Kroes, H. Y., Dempsey, J. C., Gremmels, H., Monroe, G. R., Phelps, I. G., Duran, K. J., Adkins, J., Kumar, S. A., Knutzen, D. M., Knoers, N. V., Mendelsohn, N. J., Neubauer, D., Mastroyianni, S. D., Vogt, J., Worgan, L., Karp, N., Bowdin, S., … Doherty, D. (2016). MKS1 regulates ciliary INPP5E levels in Joubert syndrome. Journal of Medical Genetics, 53 (1), 62–72.

Starich, T. A., Herman, R. K., Kari, C. K., Yeh, W. H., Schackwitz, W. S., Schuyler, M. W., Collet, J., Thomas, J. H., & Riddle, D. L. (1995). Mutations affecting the chemosensory neurons of Caenorhabditis elegans. Genetics, 139(1), 171–188.

Town, T., Breunig, J. J., Sarkisian, M. R., Spilianakis, C., Ayoub, A. E., Liu, X., Ferrandino, A. F., Gallagher, A. R., Li, M. O., Rakic, P., & Flavell, R. A. (2008). The stumpy gene is required for mammalian ciliogenesis. Proceedings of the National Academy of Sciences of the United States of America, 105(8), 2853–2858.

van Dam, T. J. P., Kennedy, J., van der Lee, R., de Vrieze, E., Wunderlich, K. A., Rix, S., Dougherty, G. W., Lambacher, N. J., Li, C., Jensen, V. L., Leroux, M. R., Hjeij, R., Horn, N., Texier, Y., Wissinger, Y., van Reeuwijk, J., Wheway, G., Knapp, B., Scheel, J. F., … Huynen, M. A. (2019). CiliaCarta: An integrated and validated compendium of ciliary genes. PloS One, 14(5), e0216705.

Wheway, G., Mitchison, H. M., & Genomics England Research Consortium. (2019). Opportunities and Challenges for Molecular Understanding of Ciliopathies-The 100,000 Genomes Project. Frontiers in Genetics, 10, 127.

Wiegering, A., Dildrop, R., Kalfhues, L., Spychala, A., Kuschel, S., Lier, J. M., Zobel, T., Dahmen, S., Leu, T., Struchtrup, A., Legendre, F., Vesque, C., Schneider-Maunoury, S., Saunier, S., Rüther, U., & Gerhardt, C. (2018). Cell type-specific regulation of ciliary transition zone assembly in vertebrates. The EMBO Journal, 37(10). https://doi.org/10.15252/embj.201797791

Williams, C. L., Li, C., Kida, K., Inglis, P. N., Mohan, S., Semenec, L., Bialas, N. J., Stupay, R. M., Chen, N., Blacque, O. E., Yoder, B. K., & Leroux, M. R. (2011). MKS and NPHP modules cooperate to establish basal body/transition zone membrane associations and ciliary gate function during ciliogenesis. The Journal of Cell Biology, 192 (6), 1023–1041.

Williams, C. L., Masyukova, S. V., & Yoder, B. K. (2010). Normal ciliogenesis requires synergy between the cystic kidney disease genes MKS-3 and NPHP-4. Journal of the American Society of Nephrology: JASN, 21(5), 782–793.

Williams, C. L., Winkelbauer, M. E., Schafer, J. C., Michaud, E. J., & Yoder, B. K. (2008). Functional redundancy of the B9 proteins and nephrocystins in Caenorhabditis elegans ciliogenesis. Molecular Biology of the Cell, 19(5), 2154–2168.

Winkelbauer, M. E., Schafer, J. C., Haycraft, C. J., Swoboda, P., & Yoder, B. K. (2005). The C. elegans homologs of nephrocystin-1 and nephrocystin-4 are cilia transition zone proteins involved in chemosensory perception. Journal of Cell Science, 118(Pt 23), 5575–5587.

Yee, L. E., Garcia-Gonzalo, F. R., Bowie, R. V., Li, C., Kennedy, J. K., Ashrafi, K., Blacque, O. E., Leroux, M. R., & Reiter, J. F. (2015). Conserved Genetic Interactions between Ciliopathy Complexes Cooperatively Support Ciliogenesis and Ciliary Signaling. PLoS Genetics, 11(11), e1005627.

Zhang, D., & Aravind, L. (2010). Identification of novel families and classification of the C2 domain superfamily elucidate the origin and evolution of membrane targeting activities in eukaryotes. Gene, 469(1-2), 18–30.

Zhao, C., & Malicki, J. (2011). Nephrocystins and MKS proteins interact with IFT particle and facilitate transport of selected ciliary cargos. The EMBO Journal, 30(13), 2532–2544.

