## Supplemental Data for "Interpreting the pathogenicity of Joubert Syndrome missense variants in *Caenorhabditis elegans*" (from another version of this preprint: version 1, 2020-05-22)

### Figure S1

#### Design of *mksr-2* repair templates

100nt ssODN repair templates were engineered to contain the desired missense allele, introduce silent mutations that will prevent Cas9 from binding/cutting the repair template, and introduce a restriction enzyme site for detection of the edit in the *C. elegans* genome.

coding mutations
silent mutations
restriction site

PAM ← guide

WT ...GTTGG**CCC**CGACTTTTGCTCCAAATCTGGCACC...
...GTTGG**T**CG**C**G**T**CTC**C**TTGCTCCAAATCTGGCACC...
P74S

guide → PAM

WT ...GCGAGAAGATTGTCAGTGTGTCAGC**TGG**AGTCGT...
...GCGAGAAGATTGTC**TC**TGT**C**TC**C**GCT**AGC**GTCGT...
G155S

#### Single injection to generate both variants and detect with restriction enzyme digest

*unc-58* co-CRISPR
P74S guide and repair template
G155S guide and repair template

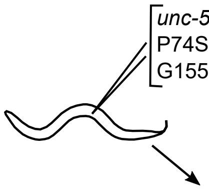

Screen F1 for dominant Unc
Make pools of F1 (2-4 worms each)
gDNA lysis
2 PCRs and 2 digests to detect variants

↓

Isolate single non-Unc F2 from F1 pools of interest
Genotype to find homozygous variants
Sequence to confirm allele

**Screening F1 progeny for P74S**

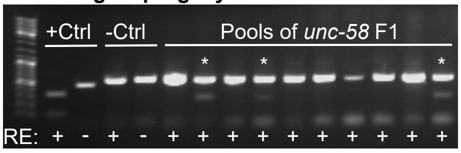


**Screening F1 progeny for G155S**

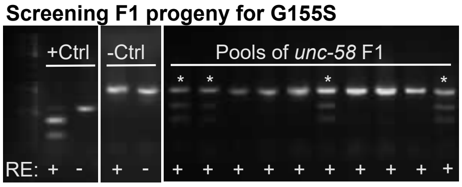


#### Simplified variant detection strategy using single PCR

Same approach to generate the edits as described above.

Detection of variants in F1 pools with single 3 primer PCR:
- Common primer
- P74S mutant primer
- G155S mutant primer

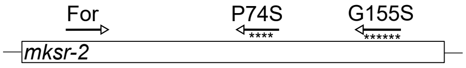


**Screening F1 progeny for both P74S and G155S**

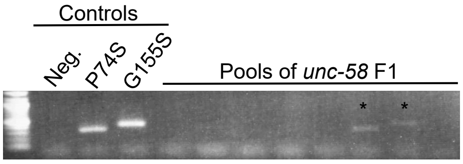


Wild-type controls will not amplify a band but the engineered alleles will produce a PCR product.

#### The simplified stragegy to identify the engineered alleles has many benefits:

- Faster screening of F1 pools (half the number of PCRs and no digests required)
- Cheaper because it requires less Taq and no restriction enzymes
- More versatile because design of alleles is not limited by restriction enzyme sequences

**Figure S1. Design and detection of pathogenic *mksr-2* variants P74S and G155S using CRISPR/Cas9**

PAM sequences targeted are shown in bold and the targeted codon is underlined. 100 nucleotide single stranded DNA repair templates were designed to introduce the desired coding variants (red), silent missense mutations (blue), and a restriction site (grey). Silent mutations were included to disrupt the sequence complementary to the guide RNA. The P74S variant introduces a BstUI site while G155S introduces an NheI site. A single injection mix containing the P74S guide RNA, P74S repair template, G155S guide RNA, G155S repair template, and *unc-58* co-CRISPR reagents was used to generate both alleles in one round of injection. 2-4 Unc F1 were pooled to screen for the engineered variants. Two PCR were performed, one for each variant, and then subsequently digested with the appropriate enzyme. The negative control gDNA was isolated from wild-type non-injected worms. The positive control is a PCR product that contains both BstUI and NheI restriction enzyme cut sites. One positive F1 pool was selected for each variant and homozygotes were isolated in the next generation. We sequenced *mksr-2* in both alleles to verify that only the desired engineered mutations were present. This approach was used to generate the mNG tagged variants *mksr-2(oq125 mNG::MKSR-2[P74S])* and *mksr-2(oq126 mNG::MKSR-2[G155S])*.

To make the non-tagged version of the variants we used a modified detection strategy. The injection strategy was not altered and these variants were also generated in a single injection. To detect the variants in the F1 pools we performed a single PCR with 3 primers: One common forward primer and two mutant reverse primers that bind the region containing the missense and silent mutations introduced by the engineered alleles. In this PCR approach the wild-type negative control does not generate a PCR product. The previously generated *mksr-2(oq125)* and *mksr-2(oq126)* strains were used as positive controls. This detection strategy was used to generate the *mksr-2(oq137[P74S])* and *mksr-2(oq138[G155S])* variants. This simplified approach to detect the variants is faster, less expensive, and more versatile.

Figure S2

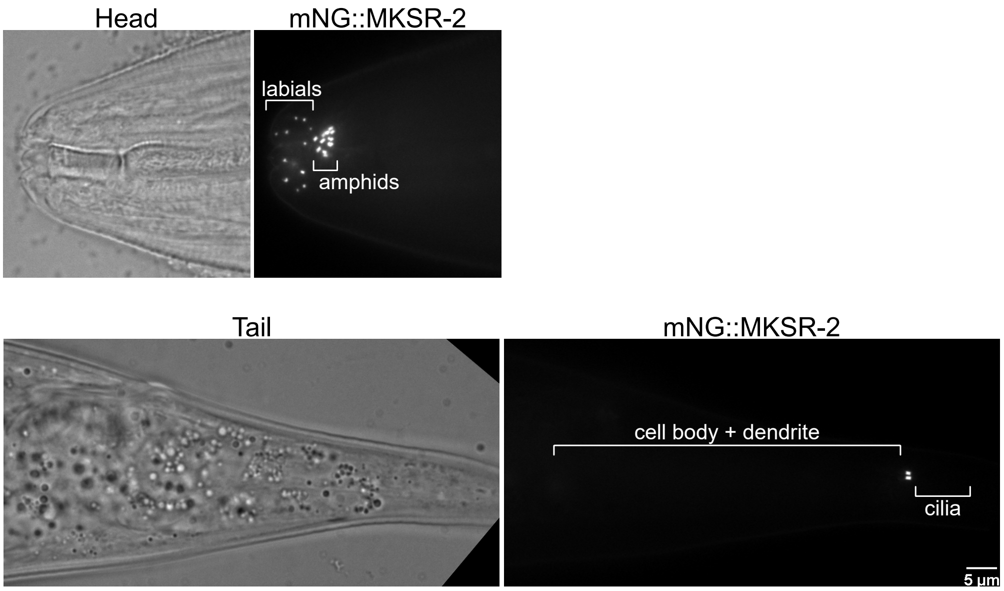


**Figure S2. (Related to Figure 2) mNG::MKSR-2 is specifically localized to the ciliary transition zone**

The fluorescence signal observed in the endogenously expressed wild-type mNG::MKSR-2 strain is very bright and specific to the transition zone. Clear transition zones can be seen in the head at the labial and amphid cilia. In the tail, mNG::MKSR-2 is observed at the transition zone in the phasmids and PQR (not pictured). Signal in the cilia, dendrite, or cell body is not observed in this strain. The scale bar is 5 µm.

### Figure S3

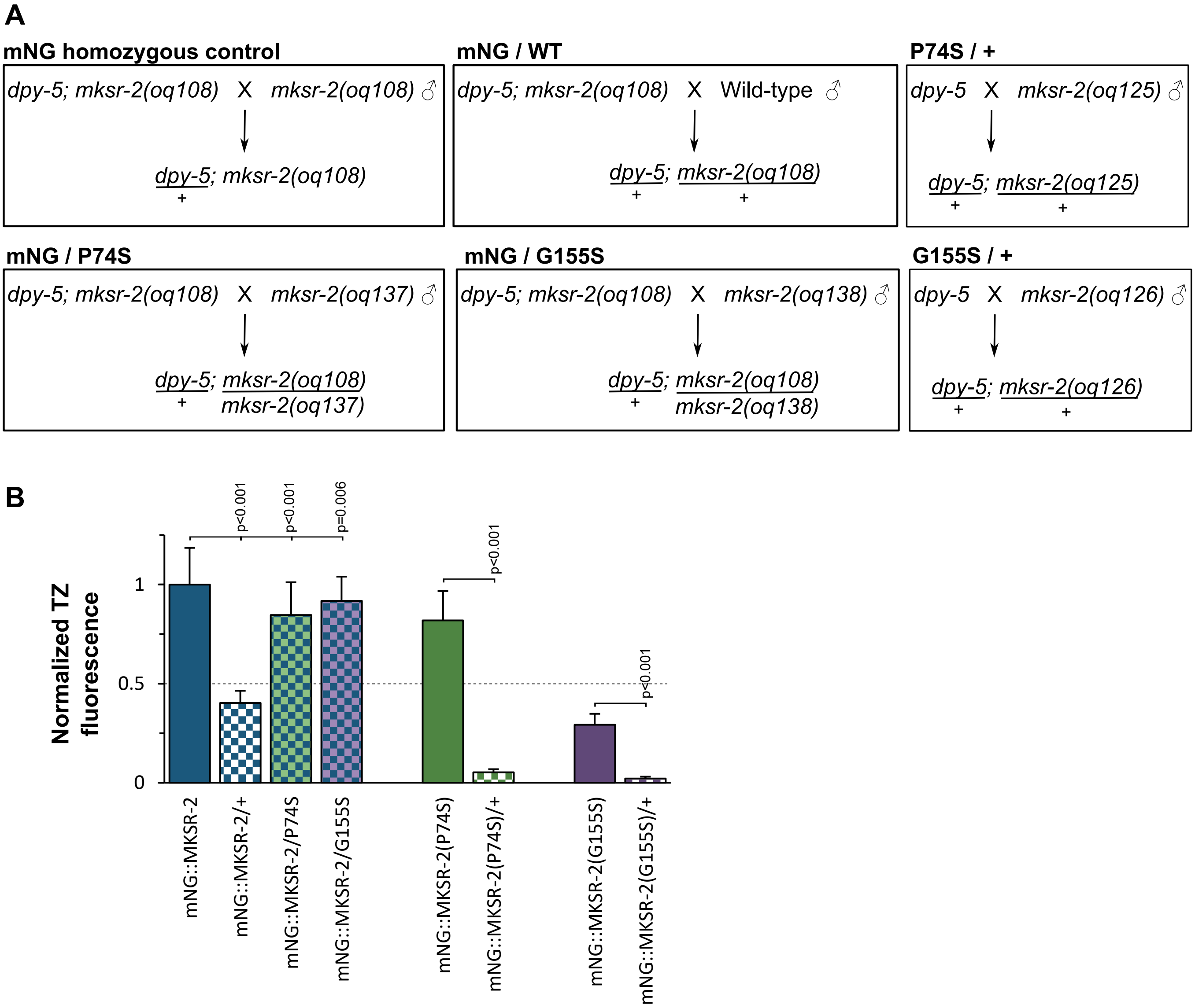


**Figure S3. (Related to Figure 2E) Quantification of fluorescence levels of heterozygous mNG::MKSR-2**

(A) Crosses used to generate F1 with mNG::MKSR-2 with a non-tagged *mksr-2* allele and to generate heterozygous F1 with mNG::MKSR-2(P74S)/+ and mNG::MKSR-2(G155S)/+ for Figure 2E.

(B) Quantification of the total fluorescence intensity of mNG::MKSR-2 at the transition zone of phasmid cilia. Background fluorescence was subtracted and the signal was normalized to homozygous mNG::MKSR-2. If both the tagged mNG and non-tagged versions of the protein are recruited to the transition zone with the same efficiency we would expect to see 50% fluorescence in the heterozygotes (grey dashed line). Error bars represent the standard deviation and all p-values were calculated in a two-tailed student's t-test. All n values are >40 phasmid pairs measured.

Figure S4

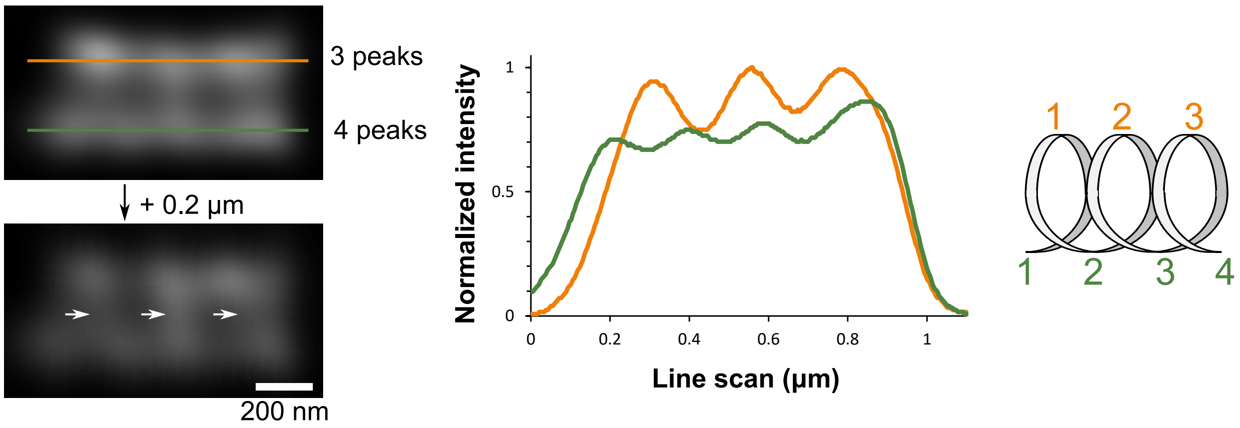


**Figure S4. (Related to Figure 2F) Confocal based super-resolution imaging of mNG::MKSR-2(+) reveals a possible spiral pattern**

The wild-type mNG::MKSR-2 fluorescence signal has discrete peaks of signal along the edge of the transition zone. In this example, line scans along the length of the transition zone show 3 and 4 discrete peaks on each side that are off set. In the next plane of the z-stack, signal that connects the opposite peaks is observed (arrows). A model whereby mNG::MKSR-2 is arranged in a spiral pattern along the transition zone could explain this observation.

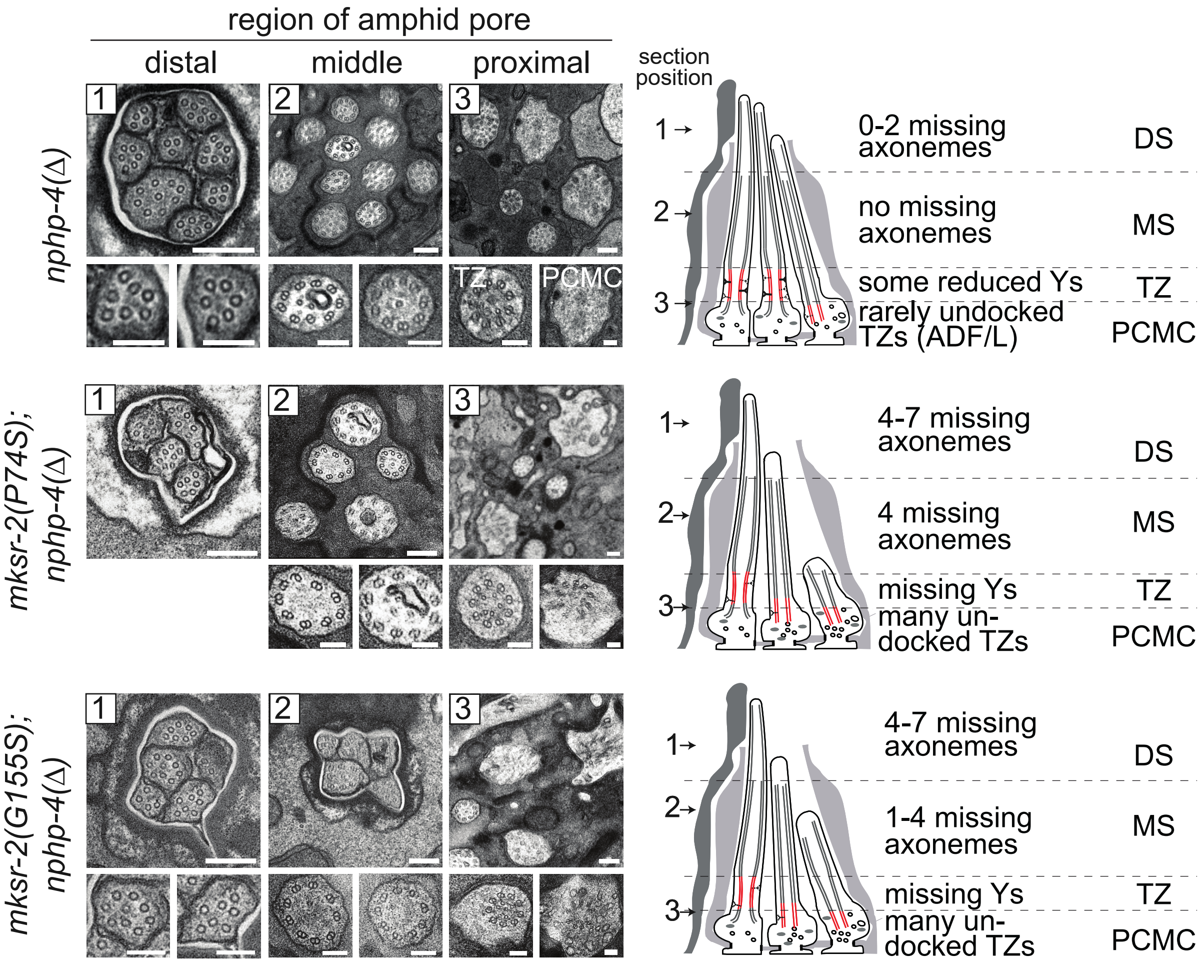


**Figure S5. (Related to Figure 4) Ultrastructure of amphid channel cilia is disrupted by P74S and G155S mutations in mksr-2.**

Transmission electron microscopy (TEM) images of the amphid sensory pore, whose channel contains 10 ciliary axonemes, in the indicated genotypes. Images are from cross sections taken from the distal (1), middle (2) and proximal (3) regions of the pore (see schematic for section positions; section position also labelled on the large images). Large images show the pore at low magnification; smaller images show individual cilia at higher magnification. Note that the *nphp-4(Δ)* control images are adapted from (Lambacher et al., 2016). Schematics of the amphid channel (only 3 of the 10 cilia are shown for simplicity) summarize the major ultrastructural features observed. DS; distal segment. MS; middle segment. TZ; transition zone. PCMC; periciliary membrane compartment. Scale bars; 100 nm (small images) and 200 nm (large images).

Figure S6

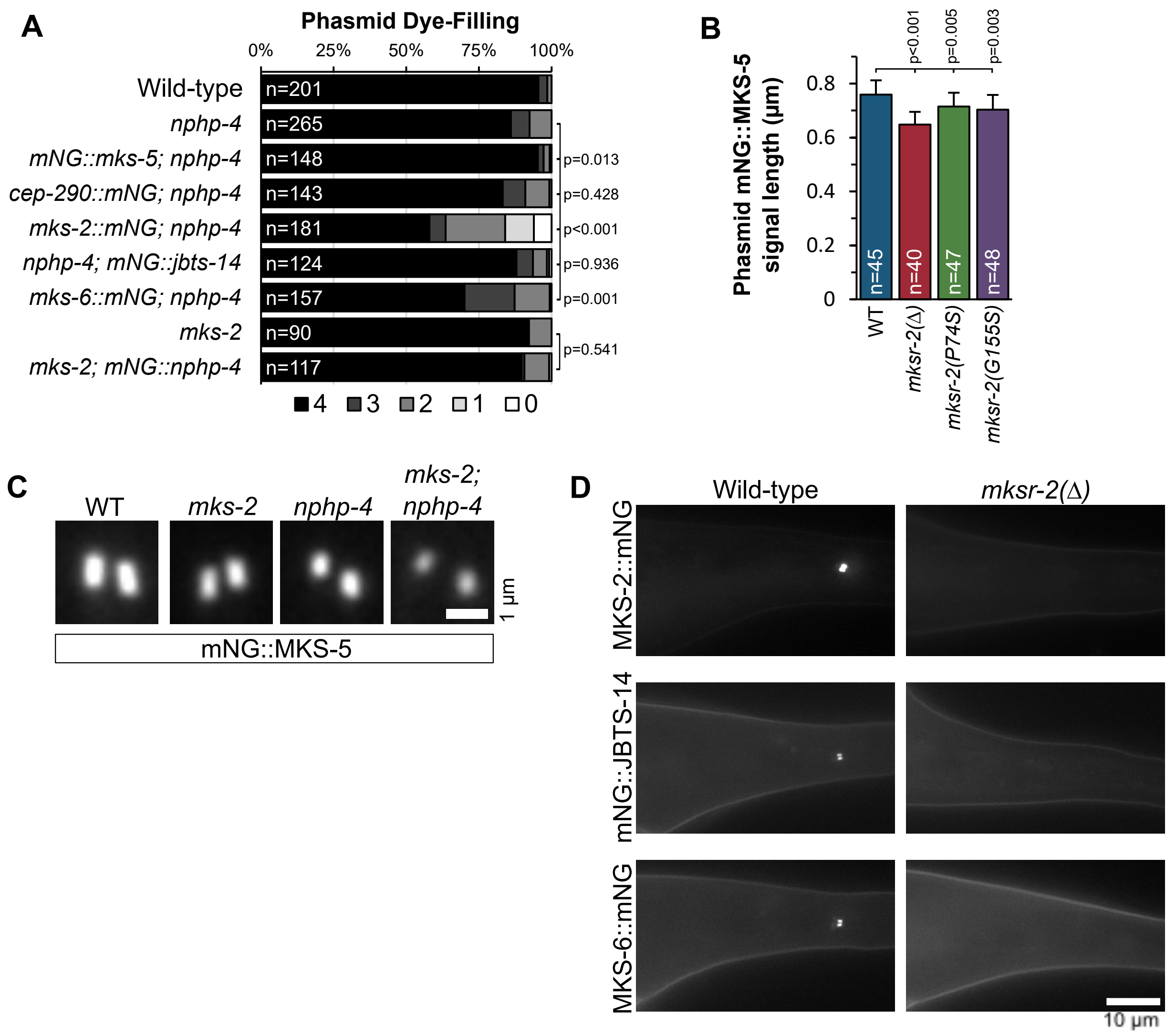


**Figure S6. (Related to Figure 6) mNG knock-ins of additional transition zone proteins**
(A) To assess if the endogenous mNG knock-ins of NPHP-4, MKS-5, CEP-290, MKS-2, JBTS-14, or MKS-6 affect the protein function in the cilia we performed and quantified a dye filling assay. Assays were performed in an *nphp-2* or *mks-2* mutant background. MKS-2::mNG had many dye filling defective worms suggesting that the cilia integrity in these strains is compromised. These strains were still used in subsequent analysis because the observed defects were mild. N values are the number of worms analyzed. (B) The length of the mNG::MKS-5 signal was measured in various *mksr-2* mutants by calculating the "full width at half max" (FWHM) from a line scan down the length of the transition zone. N value is the number of transition zones measured. All p-values were calculated in a two-tailed student's t-test. (C) mNG::MKS-5 fluorescence in wild-type, *mks-2, nphp-4,* and the double mutant. Only one punctae of signal is visible in all genotypes. The fluorescence signal was consistently fainter in the double mutant. The scale bar is 1 µm. (D) MKS-2, JBTS-14, and MKS-6 are lost from the transition zone in the *mksr-2* null mutant. The scale bar is 10 µm.

Figure S7

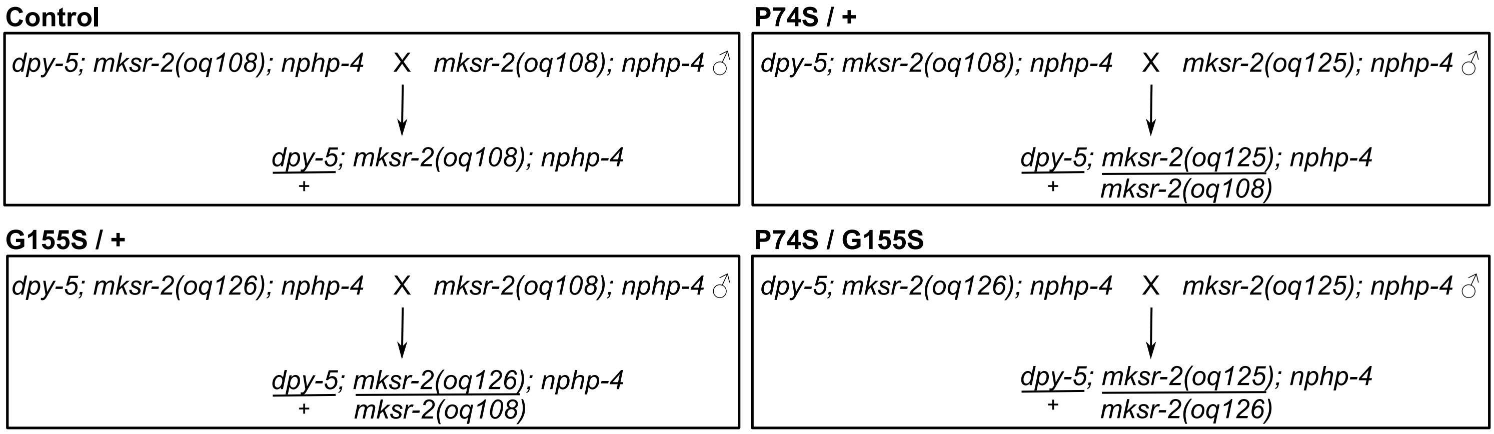


**Figure S7. (Related to Figure 7) Full genotypes of crosses used to generate heterozygous worms**
Crosses used to generate control, P74S/+, G155S/+, and P74S/G155S progeny for analysis in Figure 7 C, D, and E. All F1 progeny are homozygous for *nphp-4* and heterozygous for *dpy-5*.

### Table S1. List of CRISPR/Cas9 guides and repair templates used in this study

| Gene | | Alleles generated | Guide Sequence | Linker | Repair Template |
|---|---|---|---|---|---|
| *dpy-10* | co-CRISPR | --- | GCTACCATAGGCACCACGAG | --- | ssODN |
| *unc-58* | co-CRISPR | --- | ATCCACGCACATGGTCACTA | --- | ssODN |
| *mksr-2* | mNG N-term knock-in | *oq108* | CTTCGGCCATtttatcccgg | GTGGGGS | PCR product |
| *mksr-2* | P74S | *oq125, oq137* | CAGATTTGGAGCAAAAGTCG | --- | ssODN |
| *mksr-2* | G155S | *oq126, oq138* | AAGATTGTCAGTGTGTCAGC | --- | ssODN |
| *nphp-4* | mNG N-term knock-in | *oq109* | gaaaaATGTCGGTCAACGAC | GTGGGGSGGGGS | PCR product |
| *mks-5* | mNG N-term knock-in | *oq112* | AGGATACCTTGCGGATACCc | GTGGGGSGGGGS | PCR product |
| *cep-290* | mNG C-term knock-in | *oq103* | GTGGTAAGTGAACAGTGTTT | GTGGGGSGGGGS | PCR product |
| *mks-2* | mNG C-term knock-in | *oq101* | ttatgacgcatgTCAGCCAC | GTGGGGS | PCR product |
| *jbts-14* | mNG N-term knock-in | *oq127* | GGAACTGGACGACTTGTTGG | GTGGGGSGGGGS | PCR product |
| *mks-6* | mNG C-term knock-in | *oq128* | TGGCTGTTCTTAAACCAAAA | GTGGGGSGGGGS | PCR product |

| Gene | Repair Template Sequence |
|---|---|
| *dpy-10* | CACTTGAACTTCAATACGGCAAGATGAGAATGACTGGAAACCGTACCGCATGCGGTGCCTATGGTAGCGGAGCTTCACATGGCTTCAGACCAACAGCCTAT |
| *unc-58* | attttgtggtataaaatagccgagttaggaaacaaatttttctttcagGTTTTTCTGTCGTTACCATGTGCGTGGATCTTGCGTCCACACATCTCAAGGCGTACTT |
| *mksr-2* | tcgcggagcccattgattccctccgggataaaATG...mNG+GGAACCGGAGGTGGCGGATCT...GCCGAAGTGTTCGTTTCGGGACAAATTTTATCGGC |
| *P74S* | agacatcatgacacccccgacacatagatggaatttccagGTTGGTCGCGTCTCTTGCTCCAAATCTGGCACCATGACAACTACGGTAGACAGGAAATT |
| *G155S* | CTCAGCGCCCTGGAGGATCCGTCGATTCGCGAGAAGATTGTCTCTGTCTCCGCTAGCGTCGTCAAATTTGAATTGAATATTATTACTAAAAATTTTCAGA |
| *nphp-4* | tactaaagtttgtggtaacaaagtctcgaaaaATG...mNG+GGAACCGGAGGTGGCGGATCTGGAGGTGGCGGATCT...GTCAACGACTGGTATTCGTTGTTTCTGGCAAACCG |
| *mks-5* | tgaaacctcttataataataatacaaaattccagG...mNG+GGAACCGGAGGTGGCGGATCTGGAGGTGGCGGATCT...GTCTCCGCCAGGTATCCTATTGAAAAATGGAGCCGCCCACAGCT |
| *cep-290* | gtttttttgtttcagAAAAATCGCATTGCTCATCTtCAGAACACCGTCCACCTTCCACAA...GGAACCGGAGGTGGCGGATCTGGAGGTGGCGGATCC+mNG...TGAtttttttaataattatttttctgtaagaataa |
| *mks-2* | tttaaattttgaaaatatatttccagACAACCCGCTAGT...GGAACCGGAGGTGGCGGATCT+mNG...TGAcatgcgtcataatccttcaaatcgaattagat |
| *jbts-14* | tagatttttcacttaccctctttcttttcaagATG...mNG+GGAACCGGAGGTGGCGGATCTGGAGGTGGCGGATCT...CCCCCAACctccCGcCCAGTTCCTCCACCTCCCAAAAGAAGAATCAGgta |
| *mks-6* | CATGTTATACAAATCGATATTGCTCTGGCTGTTCTcAAgCCAAAgcGc...GGAACCGGAGGTGGCGGATCTGGAGGTGGCGGATCC+mNG...TAGaatttgaacttataactacatatatacattta |

#### Table S2. List of worms strains used in this study

| Strain | Genotype | Details |
|---|---|---|
| N2 | Wild-type | |
| | *nphp-4(tm925) V* | Deletion, null |
| | *mksr-2(tm2452) IV* | Deletion, null |
| OEB919 | *mksr-2(oq108) IV* | mNG::MKSR-2(+) |
| OEB936 | *mksr-2(oq125) IV* | mNG::MKSR-2(P74S) |
| OEB937 | *mksr-2(oq126) IV* | mNG::MKSR-2(G155S) |
| | *mksr-2(oq137) IV* | MKSR-2(P74S) |
| | *mksr-2(oq138) IV* | MKSR-2(G155S) |
| | *mksr-2(tm2452); nphp-4(tm925)* | |
| | *mksr-2(oq108); nphp-4(tm925)* | |
| | *mksr-2(oq125); nphp-4(tm925)* | |
| | *mksr-2(oq126); nphp-4(tm925)* | |
| OEB958 | *mksr-2(oq137); nphp-4(tm925)* | |
| OEB959 | *mksr-2(oq138); nphp-4(tm925)* | |
| YH746 | *yhEx414[rpi-2::gfp + osm-5p::xbx1::tdTomato + pRF4]* | |
| | *mksr-2(tm2452); yhEx414* | |
| | *mksr-2(oq108); yhEx414* | |
| | *mksr-2(oq125); yhEx414* | |
| | *mksr-2(oq126); yhEx414* | |
| OEB430 | *oqEx400[arl-13p::ARL-13::tdTomato+pRF4]* | |
| | *mksr-2(tm2452); oqEx400* | |
| | *mksr-2(oq108); oqEx400* | |
| | *mksr-2(oq125); oqEx400* | |
| | *mksr-2(oq126); oqEx400* | |
| | *mksr-2(oq108); xbx-1(cas502) V* | XBX-1::tagRFP::3xFLAG |
| | *mksr-2(oq125); xbx-1(cas502)* | |
| | *mksr-2(oq126); xbx-1(cas502)* | |
| OEB913 | *mks-2(oq101) II* | MKS-2::mNG |
| | *mks-2(oq101); mksr-2(tm2452)* | |
| | *mks-2(oq101); mksr-2(oq137)* | |
| | *mks-2(oq101); mksr-2(oq138)* | |
| OEB914 | *cep-290(oq103) I* | CEP-290::mNG |
| | *cep-290(oq103); mksr-2(tm2452)* | |
| | *cep-290(oq103); mksr-2(oq137)* | |
| | *cep-290(oq103); mksr-2(oq138)* | |
| OEB920 | *nphp-4(oq109) V* | mNG::NPHP-4 |
| | *mksr-2(tm2452); nphp-4(oq109)* | |
| | *mksr-2(oq137); nphp-4(oq109)* | |
| | *mksr-2(oq138); nphp-4(oq109)* | |
| OEB923 | *mks-5(oq112) II* | mNG::MKS-5 |
| | *mks-5(oq112); mksr-2(tm2452)* | |
| | *mks-5(oq112); mksr-2(oq137)* | |
| | *mks-5(oq112); mksr-2(oq138)* | |
| OEB938 | *jbts-14(oq127) X* | mNG::JBTS-14 |
| | *mksr-2(tm2452); jbts-14(oq127)* | |
| | *mksr-2(oq137); jbts-14(oq127)* | |
| | *mksr-2(oq138); jbts-14(oq127)* | |
| OEB939 | *mks-6(oq128) I* | MKS-6::mNG |
| | *mks-6(oq128); mksr-2(tm2452)* | |
| | *mks-6(oq128); mksr-2(oq137)* | |
| | *mks-6(oq128); mksr-2(oq138)* | |
| | *dpy-5(e907) I* | Dpy |
| | *dpy-5(e907); mksr-2(oq108)* | Dpy |
| | *dpy-5(e907); mksr-2(oq108); nphp-4(tm925)* | Dpy |
| | *dpy-5(e907); mksr-2(oq126)* | Dpy |
| | *dpy-5(e907); mksr-2(oq1026); nphp-4(tm925)* | Dpy |
